# FMRP Control of Ribosome Translocation Promotes Chromatin Modifications and Alternative Splicing of Neuronal Genes Linked to Autism

**DOI:** 10.1101/801076

**Authors:** Sneha Shah, Gemma Molinaro, Botao Liu, Ruijia Wang, Kimberly M. Huber, Joel D. Richter

## Abstract

Silencing of *FMR1* and loss of its gene product FMRP results in Fragile X Syndrome. FMRP binds brain mRNAs and inhibits polypeptide elongation. Using ribosome profiling of the hippocampus, we find that ribosome footprint levels in *Fmr1*-deficient tissue mostly reflect changes in RNA abundance. Profiling over a time course of ribosome runoff in wildtype tissue reveals a wide range of ribosome translocation rates; on many mRNAs, the ribosomes are stalled. Sucrose gradient ultracentrifugation of hippocampal slices after ribosome runoff reveals that FMRP co-sediments with stalled ribosomes; and its loss results in decline of ribosome stalling on specific mRNAs. One such mRNA encodes SETD2, a lysine methyltransferase that catalyzes H3K36me3. ChIP-Seq demonstrates that loss of FMRP alters the deployment of this epigenetic mark on chromatin. H3K36me3 is associated with alternative pre-RNA processing, which we find occurs in an FMRP-dependent manner on transcripts linked to neural function and autism spectrum disorders.

**Highlights:**

- Loss of FMRP results in decline of ribosome stalling on specific mRNAs (eg., SETD2)
- Increased SETD2 protein levels alter H3K36me3 marks in FMRP deficient hippocampus
- Genome-wide changes in mRNA alternative splicing occur in FMRP deficient hippocampus
- H3K36me3 marks and alternative splicing changes occur on transcripts linked to autism

## Introduction

The Fragile Syndrome (FXS) is characterized by intellectual disability, developmental delays, social impairment, hyper-anxiety, and other maladies. The root cause of FXS is an iteration of ∼200 or more CGG triplets in *FMR1* that induces DNA methylation and transcriptional inactivation. Loss of the *FMR1* gene product FMRP results in synaptic dysfunction and aberrant circuit formation, which produces neuro-pathological conditions during development (Santoro et al 2012). FXS is modeled in *Fmr1* knockout mice, which mimic many facets of the human disease including learning and memory deficits, repetitive disorders, and susceptibility to seizures. In these animals, neuronal communication is frequently examined at hippocampal Schaffer collateral-CA1 synapses, which exhibit exaggerated metabotropic glutamate receptor-dependent long term depression (mGluR-LTD) (Huber et al 2002). This form of synaptic plasticity normally requires protein synthesis but in *Fmr1*-deficient animals, protein synthesis is both unnecessary for mGluR-LTD and excessive in neurons, which contributes to aberrant circuit formation and other hallmarks of the syndrome (Waung and Huber 2009).

FMRP is an RNA binding protein present in most cells; in neurons, its localization at postsynaptic sites is thought to control activity-dependent synapse remodeling by regulating mRNA expression (Richter et al 2015). Considerable effort has been made to identify FMRP target mRNAs but the most rigorous involves *in vivo* crosslink and immunoprecipitation (CLIP). CLIP combined with RNA-seq has identified 842 target mRNAs in the p11-p25 mouse cortex and cerebellum (Darnell et al 2011), predominantly a single mRNA in cultured cortical neurons (Tabet et al 2016), 1610 RNAs in the p13 cortex, hippocampus, and cerebellum (Maurin et al 2018), and ∼6000 RNAs in HEK cells ectopically expressing epitope-tagged FMRP (Ascano et al 2012). This diversity of FMRP CLIP targets could reflect the different procedures employed or the different brain cell types examined. Surprisingly, the majority of CLIP sites in mRNA are in coding regions (Darnell et al 2011; Maurin et al 2018), with perhaps a bias to some cis sequences (Anderson et al 2016; Maurin et al 2018). FMRP association with mRNA coding regions and co-sedimentation with polyribosomes in sucrose gradients (Khandjian et al 1996; Feng et al 1997; Stefani et al 2004) suggests that it normally inhibits translation by impeding ribosome translocation. Indeed, studies that examined polypeptide elongation directly (Udagawa et al 2013) or susceptibility to puromycin release of nascent polypeptides (Darnell et al 2011) indicate that FMRP regulates translation at the level of ribosome transit. Supporting evidence comes from studies showing that *Drosophila* FMRP interacts with ribosomal protein L5 (rpL5) (Ishizuka et al 2002), which may preclude tRNA or elongation factors from engaging the ribosome, thereby leading to a stalled configuration (Chen et al 2014). Although these observations need to be extended to mammalian FMRP, they suggest a molecular mechanism by which FMRP stalls ribosomes. However, such observations do not indicate whether or how FMRP could stall ribosomes on specific mRNAs.

Ribosome profiling is a whole genome method for analyzing the number and positions of ribosomes associated with mRNA; when combined with RNA-seq, it yields valuable and heretofore unobtainable information on gene expression at high resolution (Ingolia et al 2009; Ingolia 2014; Brar and Weismann 2015). We have used this method to explore FMRP regulation of gene expression in mouse adult neural stems cells (aNSCs), which in FXS have an abnormal proclivity to differentiate into glia at the expense of neurons (Liu et al 2018; Luo et al 2010; Guo 2011). We identified hundreds of mRNAs whose ribosome occupancy was up or down regulated in FMRP-deficient cells, demonstrated that hundreds of other mRNAs had altered steady state levels and whose ribosome association reflected these alterations, and discovered yet additional mRNAs whose ribosome occupancy was “buffered” so that changes in their levels was compensated for by opposite changes in ribosome association (Liu et al 2018).

Ribosome profiling at steady state cannot distinguish between translocating and stalled ribosomes, suggesting that studies examining FMRP-regulated translation by this method or ribo-TRAP (Thomson et al 2017; Liu et al 2018, 2019; Greenblatt and Spradling 2018; Das Sharma et al 2019) may overlook key events leading to FXS. Moreover, if intact brain circuitry is important for FMRP regulation, then culturing disaggregated neurons, for example, might not reveal important aspects of FMRP function. To circumvent these issues, we have developed a hippocampal slice assay to detect mRNA-specific ribosome translocation dynamics. Hippocampal slices maintain the in vivo synaptic connectivity and cellular architecture, as well as express protein synthesis and FMRP-dependent forms of synaptic plasticity (Huber et al., 2002). Profiling over a time course of ribosome runoff in slices demonstrates a wide range of ribosome translocation rates on specific mRNAs, ranging from rapid (rates similar to those in ES cells, Ingolia et al 2011) to near complete stalling. A number of mRNAs retain 4-6 ribosomes after runoff in WT, but not in FMRP-deficient slices. Some RNAs associated with FMRP-stalled ribosomes encode proteins involved in neurologic function and transcriptional regulation. One such RNA encodes an epigenetic regulator, SETD2, a methyltransferase for the chromatin mark H3K36me3. We find an increase in SETD2 protein levels in *Fmr1*-deficient hippocampus. ChIP-seq demonstrates that distribution of this mark varies in a genotype (i.e, WT vs. *Fmr1*-deficient)-dependent manner in hippocampal tissue. In mammals, apart from blocking cryptic transcription initiation and DNA damage response, H3K36me3 is correlated with pre-mRNA processing events such as alternative splicing (Kim et al 2011; Kolasinska-Zwierz et al 2009). RNA sequencing reveals that a number of splicing events such as exon skipping are altered in FMRP-deficient slices, and that these transcripts are linked to neural function and autism spectrum disorders. Aberrant RNA splicing of synapse related genes has in recent years been found to be widely prevalent in autism spectrum disorders (Lee et al., 2016; Quesnel-Vallières et al., 2016; Smith and Sadee, 2011). Our study demonstrates that a reduction in ribosome stalling in the *Fmr1*-deficient brain results in a cascade of epigenetic and RNA processing changes that likely contribute to neurologic disease.

## Results

Hippocampal Schafer collateral-CA1 synapses exhibit an exaggerated mGluR-LTD in *Fmr1*-deficient mice that is aberrantly protein synthesis-independent (Nosyreva and Huber 2006; Huber et al 2000). To identify mRNAs whose basal level expression is misregulated, we prepared CA1-enriched hippocampal slices (400µm thickness) (Figure 1A), performed ribosome profiling and RNA-seq (Figure 1B) and calculated translational efficiency (TE), which is defined as the number of ribosome footprints divided by the number of RNA-seq reads. We identified RNAs with changes in TE (TE up /TE down) or RNAs whose levels were elevated or reduced but without commensurate changes in TE (mRNA up/ mRNA down) (Figure 1C, Table S1) (nominal p-value< 0.01, false discovery rate (FDR) = 0.097 by permutation test). A gene ontology (GO) functional analysis of the biological processes most enriched in each regulatory group shows that the mRNA down group encodes factors that mediate taxis/chemotaxis, while the “mRNA up” group encodes factors involved in ribonucleotide metabolism (Figure 1D, S1A, B). The “TE down” mRNAs encode factors that control synaptic transmission (Figure 1D, S1C). The “mRNA down” group is enriched for cellular components linked to the extracellular matrix while the “mRNA up” group encodes the myelin sheath and spliceosome-interacting factors. The “TE down” group is involved in the extracellular matrix and structures involved in synaptic architecture (Figure 1E, S1C). No GO terms from the TE up group were statistically significant.

**Figure 1.**
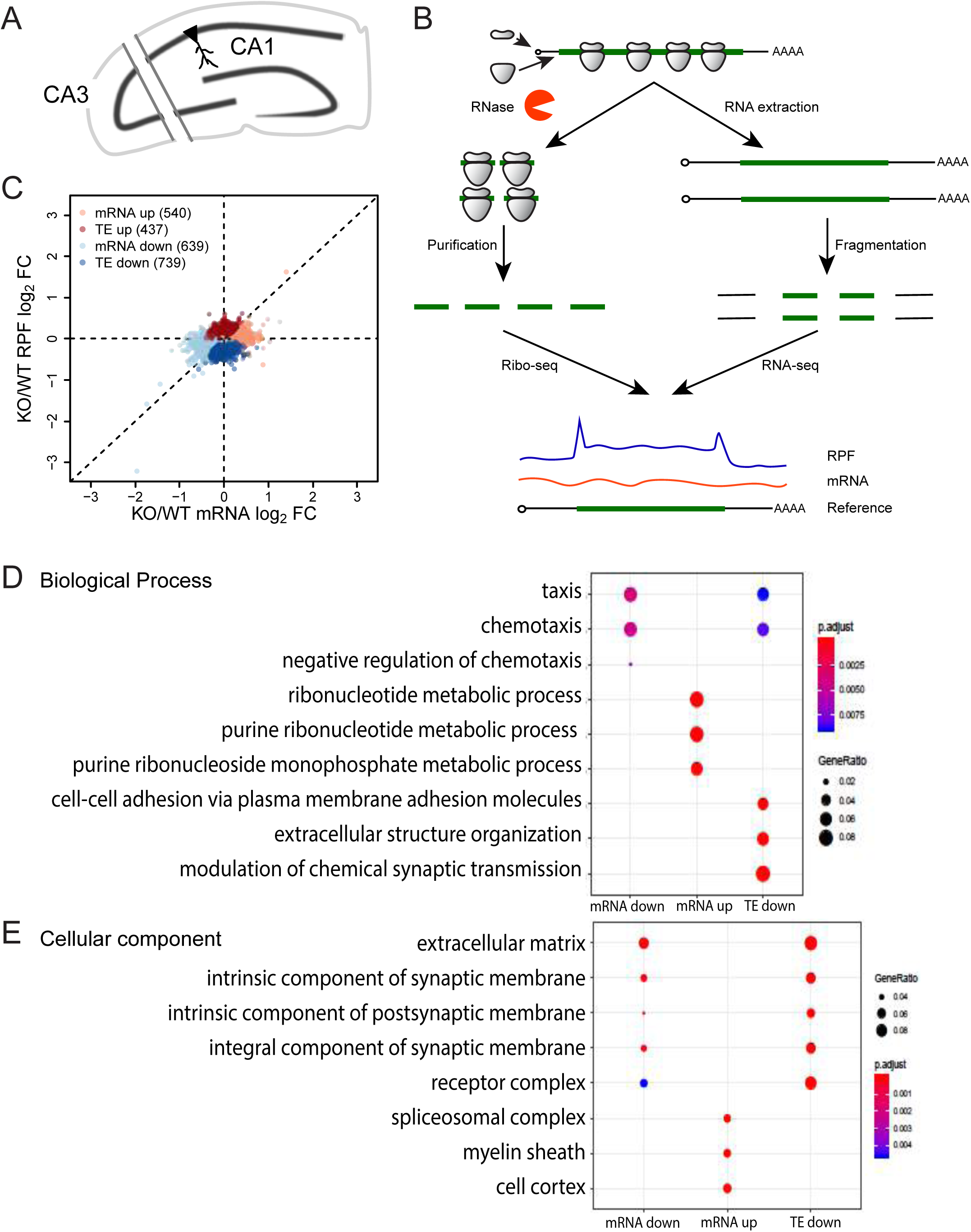
Ribosome Profiling Reveals Diverse Changes of Gene Expression in *Fmr1* KO Hippocampus. (A) Schematic diagram of the hippocampal slice preparation. To reduce spontaneous electrical activity, region CA3 was excised. (B) Schematic diagram of the experimental procedures for ribosome profiling. (C) Scatter plot of expression changes of mRNA levels and ribosome protected fragments (RPFs). Dysregulated mRNAs in the absence of FMRP are classified into four regulatory groups. 14459 genes past filtering are used for the scatter plot. Nominal p-value < 0.01, FDR = 0.097 by permutation test. FC, fold change. TE, translational efficiency. (D) Top 3 GO terms of Biological Process enriched in each regulatory group. The enrichment (gene ratio) is represented by the size of dots. The enrichment significance (adjusted p-value) is color coded. (E) Top 3 GO terms of Cellular Component enriched in each regulatory group. (also see Figure S1, Table S1)

Heat maps of the top 20 mRNAs with altered TEs both up and down in *Fmr1-*deficient hippocampal slices compared to the wild type are shown in Figures 2A and B. Examples of the change in TEs is depicted in Figure 2C. *Gfpt2* (glutamine-fructose-6-phosphate transaminase 2) mRNA, is nearly unchanged in *Fmr1* KO but has an increase in ribosome footprints Figure 2C). Conversely *Lhfpl5* (Lipoma HMGIC fusion partner-like 5) mRNA, has reduced numbers of ribosome footprints while RNA levels are mostly unchanged (Figure 2D). Neither *Gfpt2* nor *Lhfpl5* mRNAs have FMRP CLIP sites. Figure 2E shows boxplots for all 843 FMRP CLIP RNAs (p-value <0.0001, Wilcox test) (Darnell et al 2011). As a group, these RNAs decrease significantly in *Fmr1* KO, which is tracked by commensurate decreases in ribosome footprints resulting in no change in TE.

**Figure 2.**
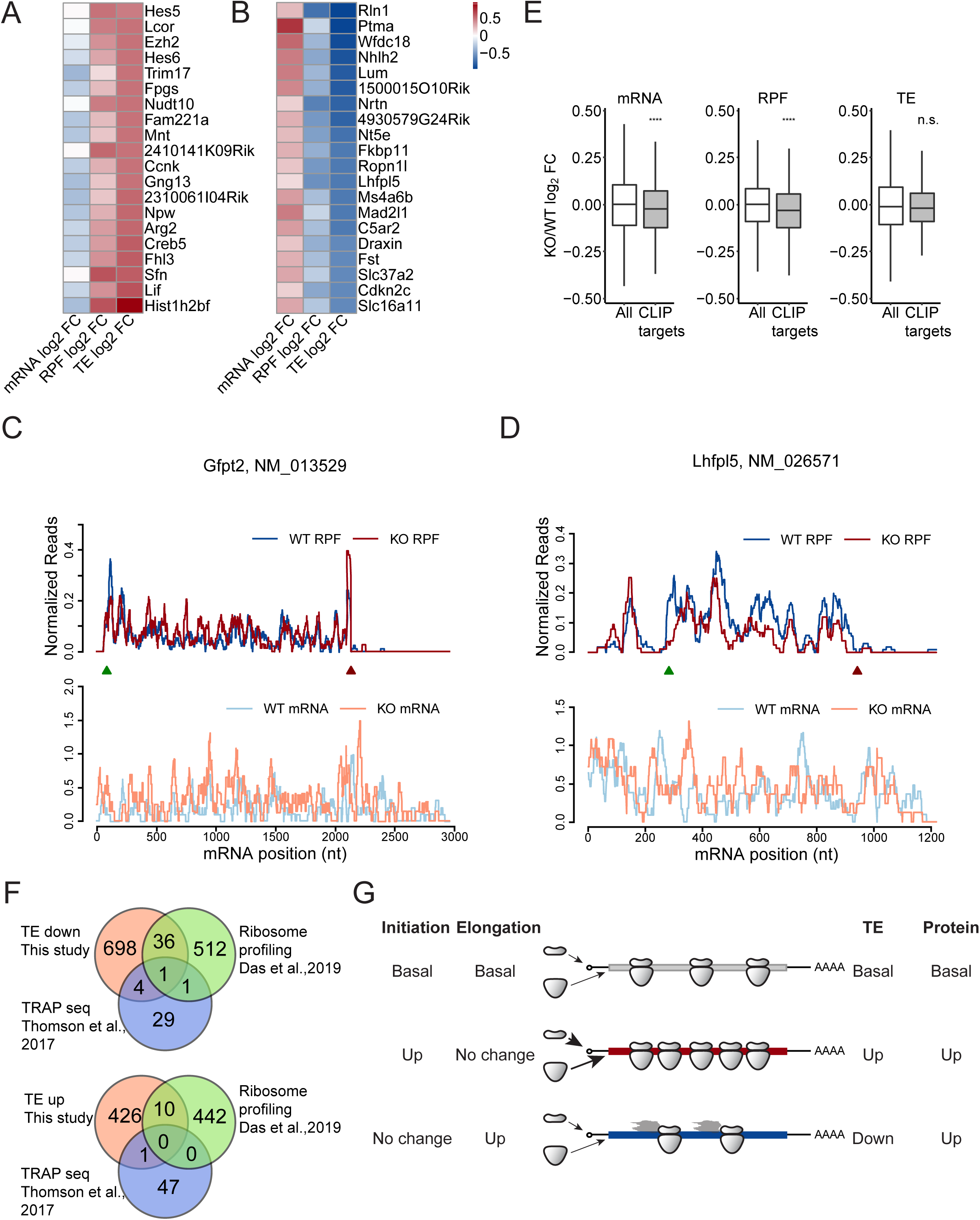
Characterization of Genes with Translational Efficiency Changes in *Fmr1* KO Hippocampus. (A) Heatmap of expression changes (log_2_FC KO/WT) for the top 20 RNAs in the “TE up” group. (B) Heatmap of expression changes (log_2_FC KO/WT) for the top 20 RNAs in the “TE down” group. (C) Read distributions on the *Gfpt2* RNA of TE up group. Normalized RPF reads (top), and mRNA reads (bottom) averaged across all replicates are plotted along the mRNA nucleotide positions with green and red triangles for annotated start and stop codons respectively. For visualization purposes, the curves were smoothed within a 30nt window. (D) Read distributions on the *Lhfpl2* RNA of TE down group. (E) Boxplots visualize the medians of expression changes for FMRP CLIP targets. The lower and upper hinges correspond to the first and third quartiles. The whiskers extend from the hinges to the largest and smallest values no further than 1.5 fold of inter-quartile range. Outliers are not shown. Gene expression changes of CLIP genes were compared to those of all genes used for DEG analysis (ns: not significant, ****, p-value <0.0001, Wilcox test). CLIP genes are the FMRP targets identified in (Darnell et al 2011). (F) Overlap of the TE down and up genes detected in this study (orange) with those of Das Sharma et al (2019 (green) and the TRAP-seq data of Thomson et al (2017) (blue). (G) Schematic models of ribosome density (TE) changes that reflect increased protein synthesis rates. (also see Table S1)

We compared our “TE down” and “TE up” groups in *Fmr1* KO hippocampus with results from Das Sharma et al (2019), who performed ribosome profiling on mouse frontal cortex, and Thomson et al (2017) who performed ribosome affinity purification (TRAP-seq) from hippocampal CA1 pyramidal neurons (Figure 2F). Only 37 of our TE down RNAs were similarly detected by Das Sharma et al (2017) (∼7% overlap) and only 5 by Thomson et al (2017) (∼ 17% overlap). Even fewer of our TE up RNAs were detected in either of those studies (Figure 2F) (∼2% overlap). We surmise that the differences in methods and source material are likely responsible for little overlap among the studies.

Our data show almost all changes in ribosome footprint number in *Fmr1*-deficient hippocampal-cortical slices can be attributed to altered RNA levels irrespective of whether the transcripts are FMRP CLIP targets. This suggests that the increase in incorporation of labeled amino acids into protein in *Fmr1* KO hippocampal slices (e.g., Dolen et al 2007; Udagawa et al 2103; Bowling et al 2019) is mostly due to changes in RNA levels and/or ribosomal transit rate (see below). However, TE may not necessarily reflect protein production. For example, increased initiation with no change in elongation would result in an increased density of ribosome footprints and yield an elevated level of protein product (Figure 2G). If there is no change in initiation but elevated elongation, the TE will appear to be reduced but the production of protein will be increased. Consequently, ribosome profiling at steady state, which cannot distinguish between transiting and stalled ribosomes, may inadequately represent translation if there are changes in elongation. To assess whether FMRP regulates ribosome dynamics, we modified our procedure to investigate temporal changes in genome-wide ribosome translocation rates.

### Ribosome run off dynamics in hippocampal-cortical slices

We next performed ribosomal profiling and RNA-seq in hippocampal-cortical slices at different time points after blocking initiation with homoharringtonine (HHT). HHT halts ribosomes on initiation codons but allows elongating ribosomes to continue translocating until they run off the mRNA (Ingolia et al 2011; Lee et al 2012). Using this assay, we aimed to determine differences in ribosome translocation rates of all expressed genes in wild type tissue slices. Slices were incubated with HHT for 0-60 min and ribosome transit stopped at specific times with cycloheximide; the samples were then used for ribosome profiling (Figure 3A, Table S1). A metagene analysis of the ribosome footprints on 1401 mRNAs with open reading frames of >3000 nt aligned with the start and stop codons is shown in Figure 3B, indicating that the ribosomes runoff in a 5’-3’ direction. We chose long mRNAs with ORFs > 3000nt to ensure that ribosome run-off was not complete within 10 min of HHT treatment and read densities could be obtained at each nucleotide position. These read densities were normalized to the average densities of the last 500 nucleotides of the CDS and is based on the P site of the ribosome footprints. Using these data, a linear regression analysis between the HHT treatment time and ribosome runoff distances was used to estimate the global elongation rate of 4.2 nt/sec at 30°C (Figure 3C). By comparison, the global elongation rate in cultured ES calls is 1.7 nt/sec at 37°C (Ingolia et al 2011).

**Figure 3.**
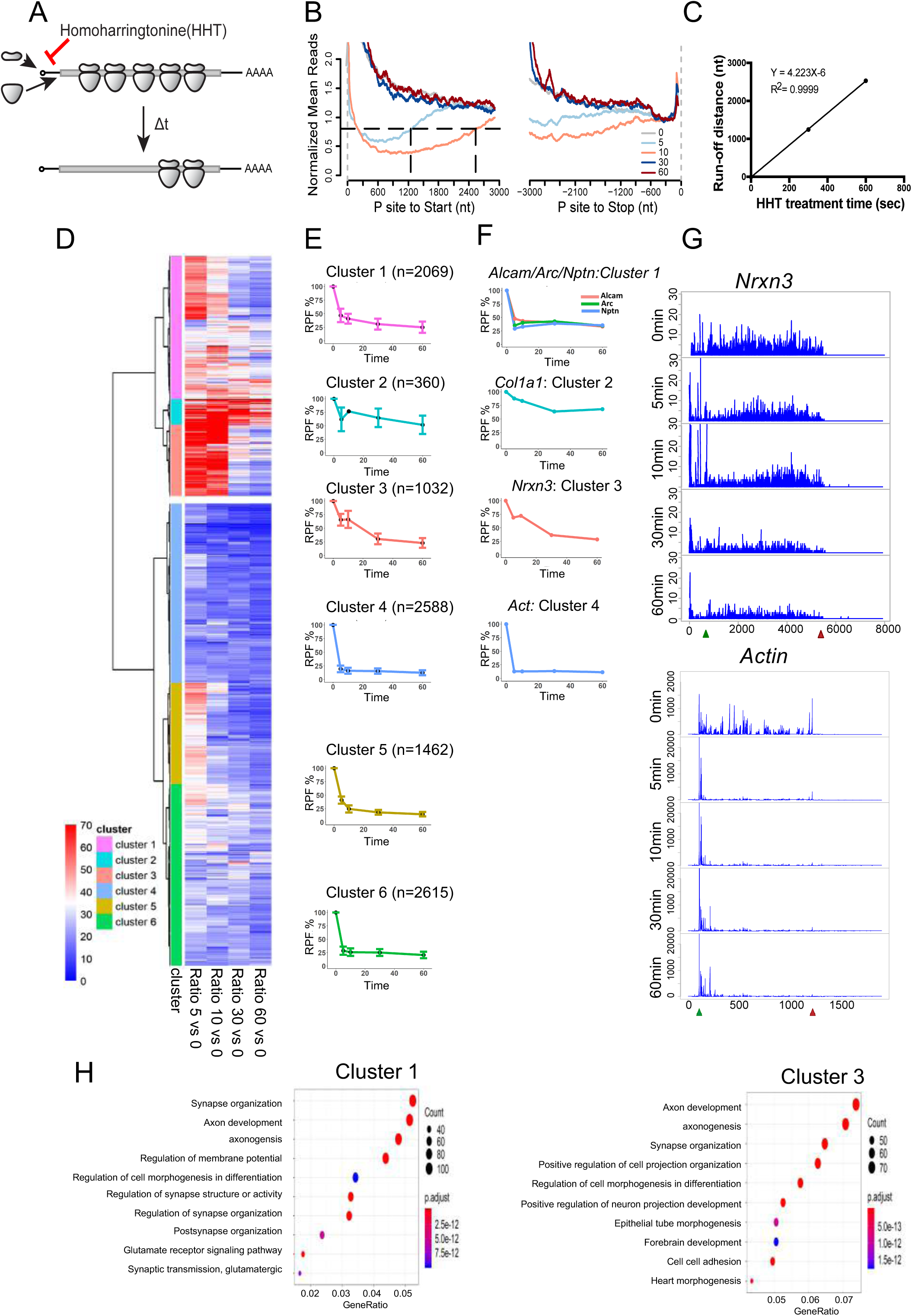
Run-off ribosome profiling of WT mouse brain slices. (A) Schematic diagram of homoharringtonine (HHT) run-off ribosome profiling. WT mouse brain splices was treated with 20µM HHT, an inhibitor of translation initiation, to allow ribosome run-off for 5, 10, 30, and 60min (t) at 30°C. (B) Metagene plot of RPFs after HHT treatment. Reads are mapped transcripts (N=1401) with CDS longer than 3000nt and aligned at the annotated start and stop codons (grey vertical dash lines). The read densities at each nucleotide position are normalized to the average density of the last 500nt of CDS and averaged using the P sites of RPFs. For visualization purposes, the curves were smoothed within a 90nt window. Black horizontal dash line indicates the arbitrary 0.8 threshold to estimate the relative run-off distances (black vertical dash lines). (C) Linear regression between the HHT treatment time and ribosome run-off distances (from B) to estimate the global elongation rate (4.2 nt/sec). (D) Cluster analysis of gene groups with distinct ribosome run-off patterns. The RPFs of each gene at each time point was normalized to time 0. The Euclidean distance matrix was then calculated, followed by hierarchical clustering using Ward’s agglomeration method (Ward, 1963). (E) Ribosome runoff patterns for each sub-cluster. The global pattern of each sub-cluster was summarized using the corresponding median and standard deviation in each timepoint. The number of RNAs in each sub-cluster is shown in parentheses. (F) Representative ribosome runoff profiles that reflect each sub-cluster. The runoff pattern for *Actin* is similar to sub-clusters 4-6. (G) Ribosome footprints for *Nrxn3* and *Actin* mRNAs during the runoff time period. (H) GO terms for sub-clusters 1 and 3. Gene ratio refers to the percentage of total differentially expressed genes in the given GO term. (also see Figure S2, Table S1)

We next performed a cluster analysis of all the mRNAs with respect to number of ribosome protected fragments at each time point (5, 10, 30, 60 min) relative to time 0. Briefly, euclidean distance was first computed, followed by hierarchical clustering using Ward’s algorithm (Ward, 1963), and the Analysis of group similarities (ANOSIM) test (Clarke, 1993) was then used to assess whether distances between clusters were statistically greater than within clusters (Figure 3D). The mRNAs fell into two broad clusters (ANOSIM R=0.56, p-value<0.001) those whose ribosome runoff occurred relatively quickly (i.e., within 5-10 min, bottom cluster), and those whose runoff occurred slowly, if at all (top cluster). These two clusters were then further grouped into 3 sub-clusters each for 6 total sub-clusters (ANOSIM R=0.64, p-value<0.001). Sub-clusters 1-3 had slow runoff times while sub-clusters 4-6 had relatively fast ribosome runoff times (Figure 3D and 3E). The ribosomes on mRNAs in sub-cluster 2 had the slowest runoff time. We next determined the ribosome runoff patterns for the mRNAs in each sub-cluster (Figure 3E). For example, mRNAs for Alcam (activated leukocyte cell adhesion molecule), Arc (activity-regulated cytoskeleton), and Nptn (neuroplastin) are present in the sub-cluster 1, Col11 (collectin subfamily 11) in sub-cluster 2, Nrxn3 (neurexin 3) in sub-cluster 3 and actin in sub-cluster 4 (Figure 3F). As examples, Figure 3G shows the ribosome footprints of Nrxn3 and actin over time; Figure S2A shows ribosome footprints for additional mRNAs. Although there was some CDS length dependency of the ribosome runoff rates, the correlation was not linear (Figure S2B). GO term analysis showed that specific biological activities were remarkably associated with each sub-cluster. Genes in sub-clusters 1 and 3 were closely associated with neural function such as axon development and synapse organization (Figure 3H). Sub-cluster 2, which contained ribosomes with the slowest runoff rates, had genes encoding the extracellular matrix, sub-cluster 4 contained genes involved in metabolic functions, sub-cluster 5 was associated with mRNA processing and regulation, and sub-cluster 6 with organelle localization, among other functions (Figure S2C). These results suggest that clusters of mRNAs with specific functions have differential ribosomal transit rates.

### FMRP controls ribosome translocation on specific mRNAs

To investigate whether FMRP controls ribosome stalling on specific mRNAs, we first determined whether it is present in ribosome-containing complexes, as suggested by earlier studies (Feng et al 1998; Khandjian et al 2004; Stefani et al 2004; El Fatimy et al 2016). Extracts from cortices were treated with nonionic detergent (NP40) in the presence or absence of RNase and sedimented through sucrose gradients. Figure S3A shows FMRP sedimented to a “heavy” ribosome-containing region of the gradient even in the presence of NP40 or RNase (Stefani et al 2004; Darnell et al 2011). These data suggest that FMRP is associated with ribosomes in heavy fractions and thus could play a role in ribosome stalling.

We next performed sucrose gradient analysis of hippocampal slices treated with HHT for 30 min (Figure 4), which allows ribosomes on most genes to run off (as shown in Figure 3). Figure 4A shows that most FMRP co-sediments with polysomes as has been shown previously (e.g., Feng et al 1998; Stefani et al 2004), but after HHT treatment, it is more prevalent in “medium” gradient fractions that correspond to 4-6 ribosomes most likely due to association with stalled ribosomes. Other proteins (e.g., UPF1, MAP2, eEF2) do not show a similar change in sedimentation (Figure S3B). A small amount of FMRP remains in a “heavy” (>7 ribosomes) region of the gradient (Figure 4B). The same experiments were performed with *Fmr1* KO slices (Figure 4C, D). RNA-seq of the “heavy” fractions showed that 1574 RNAs and 686 RNAs were reduced or elevated, respectively, in both genotypes, indicating a sensitivity or resistance to HHT treatment (Figure 4E). *Actin* is one such RNA that was sensitive to HHT that was depleted by ribosome runoff (Figure 4F), which is consistent with results from the HHT time course (Figure 3F, G). *Map1b* RNA, on the other hand, was enriched in the heavy polysomes by HHT treatment (Figure 4G). Figure 4H shows GO terms for the RNAs depleted in the heavy fractions after runoff, which include RNA processing and other factors involved in RNA metabolism. Figure 4I shows the GO terms for RNAs enriched in the heavy polysomes after HHT treatment, which include factors involved in membrane and ion transport.

**Figure 4.**
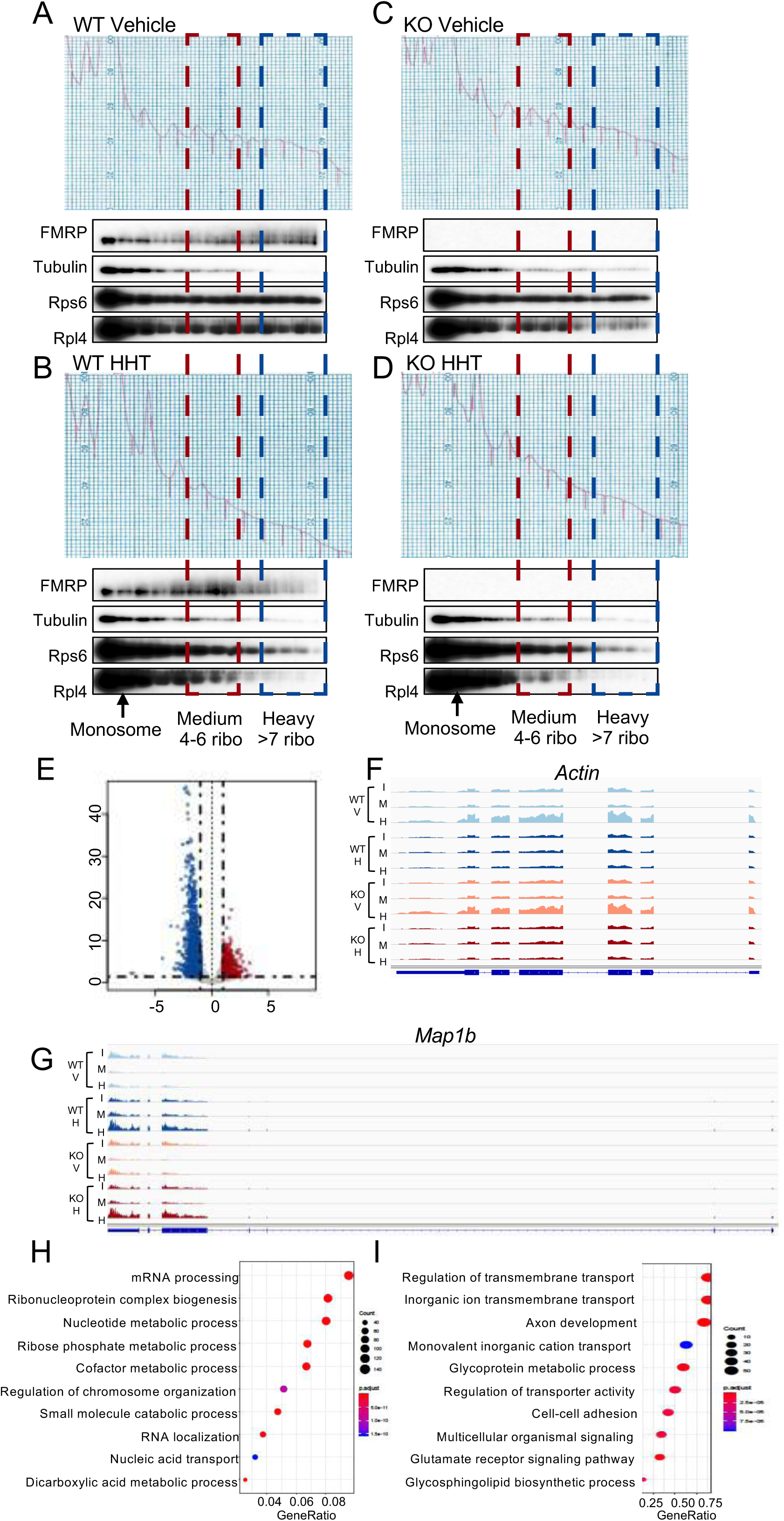
Substantial FMRP remains associated with polysomes after HHT treatment. (A-D) WT mouse hippocampal slices were treated with DMSO vehicle (A) or 20µM HHT (B) for 30min at 30°C. In parallel, *Fmr1* KO mouse hippocampal slices were also treated with DMSO vehicle (C) or 20µM HHT (D) for 30min at 30°C. Slices were homogenized and applied to 15-45% (w/w) sucrose gradients, which were fractionated with continuous monitoring of A_260_ after the ultracentrifugation. Fractions were collected for immunoblotting with indicated antibodies to detect the association of FMRP with polysomes. (E) RNA sedimenting to “heavy” fractions containing >7 ribosomes from WT slices treated with HHT was analyzed relative to input. 1574 RNAs were reduced and 686 were elevated relative to input after HHT treatment. (F) *Actin* mRNA reads in the designated conditions (WT, wild type; V, vehicle, I, input; H, heavy fractions, KO, *Fmr1* KO). (G) *Map 1b* mRNA reads in the designated conditions. (H) GO terms of RNAs depleted in heavy fractions relative to WT. (I) GO terms of RNAs enriched in heavy fractions relative to WT. (also see Figure S3, Table S1)

In contrast to the “heavy” polysome fractions, we found genotype-specific differences in the RNA-seq of the “medium” polysome fractions (Table S1). Forty-six RNAs were down-regulated in HHT-treated *Fmr1* KO slices relative to WT, indicating that they are normally bound by FMRP-stalled ribosomes (Figure 5A). Eight of these mRNAs encode epigenetic and transcriptional regulators and 23 encode proteins specifically involved in neural function (Figure 5B). One example is *Ankrd12* (ankyrin repeat domain containing 12), which is depleted in the KO slices following HHT treatment (Figure 5C). Figure 5C shows that after HHT treatment, *Ankrd12* RNA contains fewer stalled ribosomes in the medium fractions of *Fmr1* KO slices, confirming this mRNA is under FMRP regulation. Figure 5D demonstrates that compared to all RNAs, the HHT-sensitive RNAs (Figure 5B) showed significant decreases in RPFs and TEs but not RNA levels in *Fmr1* KO relative to WT (ribosome profiling data of these RNAs from Figure 2, **,p-value <0.01; ***,p-value <0.001, Wilcox test) showing the consistency between steady state ribosome profiling and the dynamic ribosome run-off assay.

**Figure 5.**
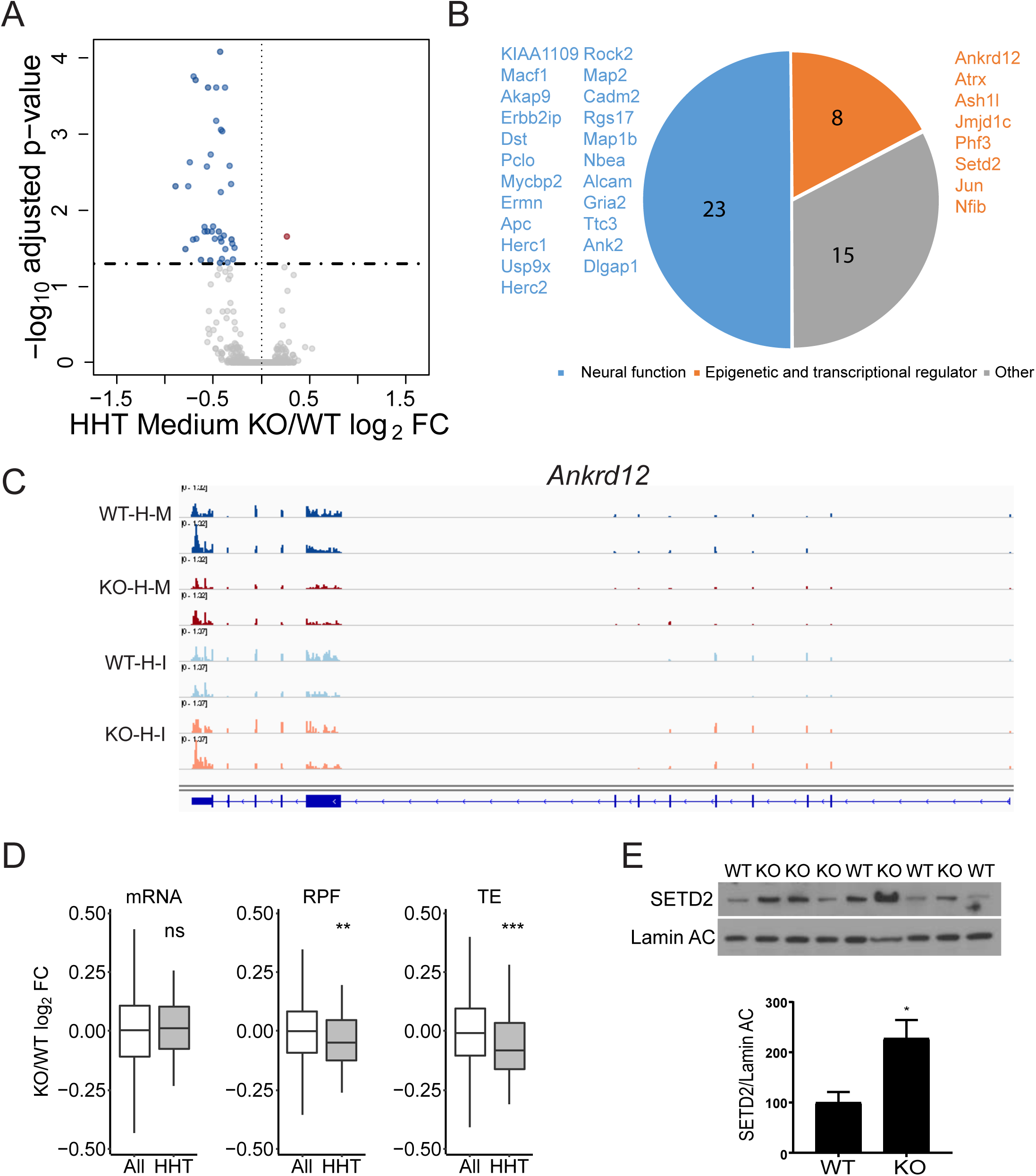
FMRP stalls ribosomes on specific mRNAs. (A) RNA sedimenting to “medium” polysomes containing 4-6 ribosomes after HHT treatment of hippocampal slices; 46 RNAs are down-regulated and 1 is up-regulated in *Fmr1* KO relative to WT. (B) Down-regulated RNAs in HHT-treated *Fmr1* KO slices primarily encode epigenetic and transcriptional regulators and proteins involved in neural function. (C) Example of *Ankrd12* RNA, which has reduced reads in *Fmr1* KO slices relative to WT after HHT (H) treatment. Input (I) reads are similar in both genotypes. M refers to medium fraction. (D) Box plot showing the fold change of *Fmr*1 KO versus WT of all RNAs (white) compared to those identified in Figure 5 A and B (gray) with respect to steady state RNA levels, RPFs, and TE (ns: not significant, **,p-value <0.01; ***,p-value <0.001, Wilcox test). (E) Western analysis of SETD2 and lamin ac in hippocampus from 4 WT and 5 *Fmr1* KO mice. When quantified and made relative to Lamin AC, SETD2 was significantly increased in the KO (p=0.0245, two-tailed t test). (also see Table S1, Figure S5 E)

Finally, Figure 5E demonstrates that one of the mRNAs decreased in the “medium” fraction in the *Fmr1* KO slices, *SETD2*, shows an increase in its protein levels in *Fmr1* KO hippocampus relative to WT (p-value=0.0245, two-tailed t test), which is expected if FMRP stalls ribosome translocation on this RNA. We find no statistically significant increase in SETD2 mRNA levels in the RNA-seq data from the FMRP KO hippocampal tissue slices (from Figure 1). We tested SETD2 protein levels in immortalized MEFs derived from WT and *Fmr1* KO mice and found no change relative to genotype (Figure S5E). This result might reflect a possible brain-specific function of FMRP or that rapidly dividing MEF cell lines do not recapitulate FMRP-dependent ribosome stalling, at least not on SETD2 mRNA. SETD2 is the main histone lysine methyltransferase that trimethylates the histone H3 tail on the lysine36(K36) residue.

### FMRP mediated regulation of H3K36me3

To investigate whether altered SETD2 levels influence gene-specific changes in H3K36me3, we performed ChIP-seq from WT and *Fmr1* KO hippocampus in duplicate, combining hippocampal tissue from 2 animals per biologic replicate (Figure 6A). All genotypic replicates were highly correlated with each other (Figure S4A). H3K36me3 is a broad histone mark that is enriched primarily in the body of the gene. To measure regions with enrichment of the mark we used the SICER v1.1 package (Xu et al., 2014) that annotates ChIP signal over a stretch of DNA as ‘islands’ (an island is defined as a cluster of non-overlapping consecutive DNA regions (200 bp windows) separated by at least a 600bp gap region of DNA with no ChIP-enriched signal detected above threshold). Annotation of the H3K36me3 islands shows that they were primarily present within the gene body compared to promoter proximal regions, as expected (Krogan et al 2003; Tiedemann et al 2016) (Figure 6B). Metagene analysis showed there to be no genotype-specific difference in the distribution of marks overall along the gene body and no effect of gene length (Figure S4B, C, D). We further compared the differential enrichment of the H3K36me3 islands within the replicates by a negative binomial test performed using edgeR (Robinson et al., 2009) (fold change >2 and p-value < 0.05). We found 363 islands with decreased and 2223 islands with increased H3K36me3 in the *Fmr1* KO compared to wild type hippocampus ChIP-seq in both replicates (Table S2). Examples of the changes of these marks that are genotype-dependent are shown in Figure 6C where H3K36me3 on *Reep4* (receptor expression-enhancing protein 4) is reduced but elevated on *Tprkb* (TP53-related protein kinase binding protein). The decreased islands were fewer and mostly found in intragenic regions whereas the increased islands were enriched and present in both the intragenic and intergenic regions (Figure 6D), demonstrating a redistribution of H3K36me3 in FXS model mice. Many of the islands, especially those that increase in *Fmr1* KO animals are found on genes linked to autism spectrum disorders as defined by SFARI (Figure 6E). A subset of genes (37) showed both increased and decreased islands, but at different locations on the gene body (Figure 6E). GO term enrichment analysis of the genes that increased H3K36me3 islands is shown in Figure 6F. No significant GO terms were enriched in the group of genes with decreased H3K36me3 islands. The genes with increased H3K36me3 islands are remarkably enriched for those involved in neural function such as synapse assembly and animal behavior. We find no correlation between steady-state RNA abundance and H3K36me3 levels on islands in the genes (Supplemental Figure S4F), which could be attributed to compensatory post-transcriptional mechanisms affecting steady state RNA. Furthermore, the methylated islands also undergo genotype-specific length changes; they are significantly longer in the WT compared to *Fmr1* KO (p-value <0.001, Kolmogorov-Smirnov (K-S) test) (Figure S4E). Our data thus show that an increase in the levels of SETD2 in the *Fmr1* KO resulting primarily in increased H3K36me3 chromatin marks at a subset of genes.

**Figure 6.**
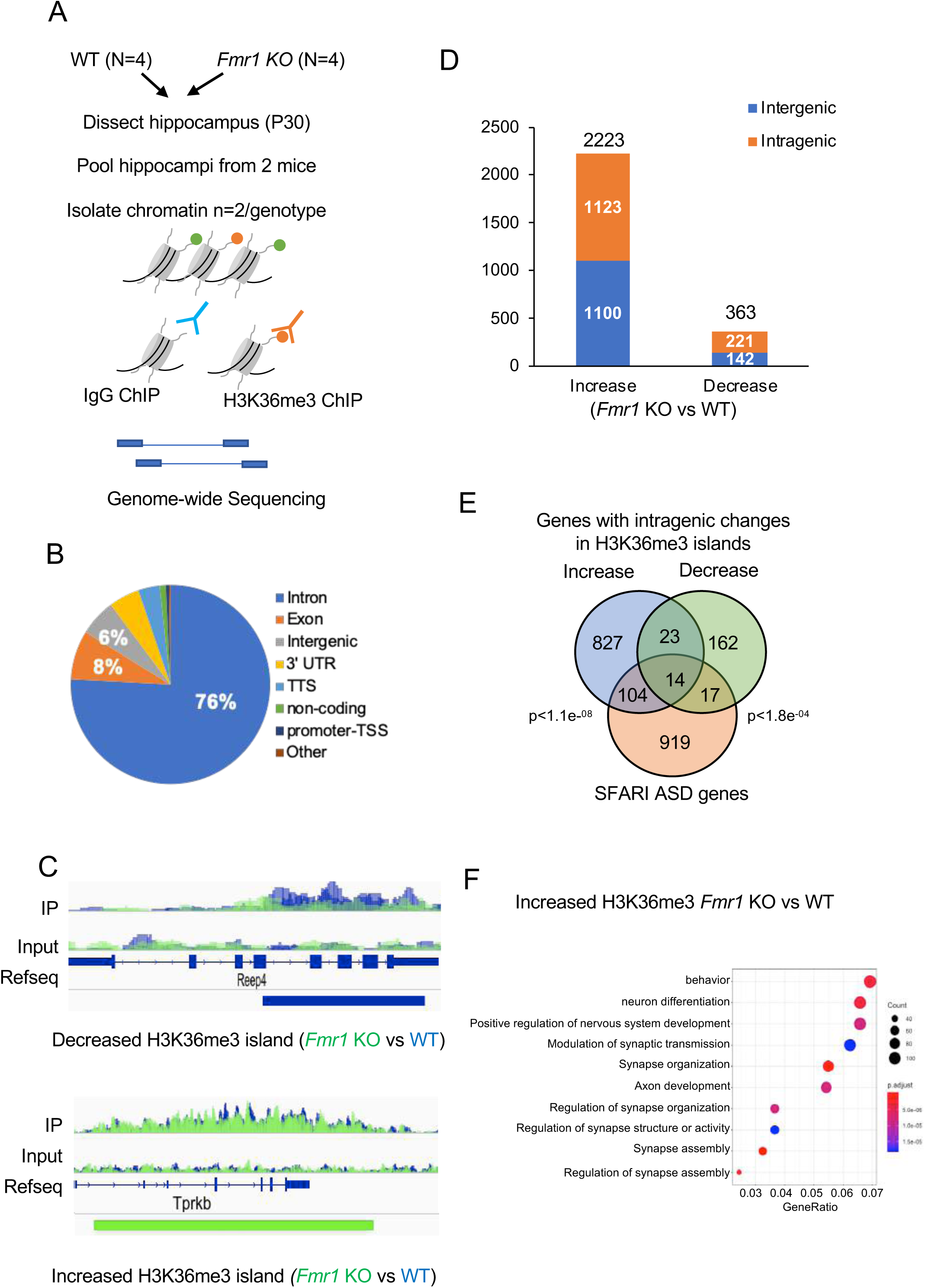
H3K36me3 localization is altered in *Fmr1* KO hippocampus. (A) Experimental design for *in vivo* ChIP-seq of H3K36me3 in hippocampus from adult WT (N=4) and *Fmr1* KO (N=4) mice. (B) Pie chart representing the genomic annotation of the total H3K36me3 islands identified in the WT and *Fmr1* KO ChIP-seq. (C) H3K36me3 ChIP-seq gene tracks for WT and *Fmr1* KO hippocampal tissue. The two sequencing tracks from each biological replicate (n=2) of WT and *Fmr1* KO were merged and overlaid. WT ChIP-seq tracks are in blue and KO tracks are in green. The tracks for IP and input are displayed. The island with significantly decreased (blue) or increased (green) tracks are shown below the Refseq gene annotation (FDR <0.0001 and p-value <0.01) as identified using the SICER package. *Reep4* shows decreased H3K36me3 islands in *Fmr1* KO and *Tprkb* shows increased islands in *Fmr1* KO (D) Distribution of significantly increased or decreased H3K36me3 islands in intragenic and intergenic regions of the genome using negative binomial test performed using edgeR (Robinson et al., 2009) (fold change >2 and p-value < 0.05). The total number of increased or decreased islands is indicated above the respective bars in the graph in *Fmr1* KO vs wild type ChIP-seq. (E) Venn diagram for significant overlap of *Fmr1* KO mis-regulated H3K36me3 genes (increased islands in blue and decreased islands in green) with the ASD-linked genes from the SFARI database. A subset of genes showed both increased and decreased islands along the length of the gene body. The p-value (hypergeometric test) is indicated next to the respective comparisons. (F) GO term enrichment of H3K36me3 differentially enriched genes in *Fmr1* KO vs WT (p. adjust value<0.05). (also see Figure S4, Table S2)

### FMRP regulates alternative RNA processing

H3K36me3 in mammals has been correlated with alternative pre-mRNA splicing (Kim et al 2011). We analyzed our RNA-seq data from WT and *Fmr1* KO hippocampal slices using rMATS alternative splicing package (Shen et al., 2014) and detected many alternative pre-mRNA processing events including skipped exon (SE), mutually excluded exon (MXE) and alternative 5’ and 3’ splice site (A5SS and A3SS) (Figure 7A, Table S3, FDR < 5% and p-value<0.05). The number of exon inclusion events are shown (Differences in exon inclusion levels is calculated as Percent Spliced In (PSI) and a cut-off of |deltaPSI| ≥ 5% is used). Several of the exon skipping events were validated by RT-qPCR, with ∼20-40% reduction in exon skipping in the *Fmr1* KO relative to WT (Figure 7B). These skipped exons have previously been found to be associated with defects in brain development and disease pathology (An and Grabowski, 2007; Fugier et al., 2011; Rockenstein et al., 1995; Wang et al., 2018). In particular, an increase in microexon 4 skipping in *Cpeb4* mRNA was found in postmortem cortical tissue from autism patients. Consequently a mouse model of *Cpeb4* microexon 4 deletion recapitulated the molecular and behavioral phenotypes found in autism patients (Irimia et al., 2014; Parras et al., 2018; Quesnel-Vallières et al., 2016). Figure 7C shows all the RNA processing events that undergo changes in *Fmr1*-deficient hippocampus relative to WT (p-value<0.05, |deltaPSI| ≥ 5%). Skipped exons are by far the most prevalent category, followed by alternative 3’ splice sites. The genes whose RNA processing events are altered in *Fmr1* KO are strongly linked to neural function such as vesicle transport to synapses and vesicle recycling (Figure 7D). Moreover, many of the alternatively spliced (Alt. Spl. or AS) genes in *Fmr1* KO hippocampus (Alt. Spl. *Fmr1* KO) are linked to autism (i.e., SFARI autism spectrum disorder database, ASD) (Figure 7E). In addition, the AS genes in *Fmr1* KO significantly overlap with genes previously found to be alternatively spliced in patients with autism (Alt. Spl. ASD) (Parikshak et al., 2016) (Figure 7E). We also find an overlap of genes with changes in H3K36me3 islands with those found to harbor aberrant splicing in *Fmr1* KO tissue (Figure 7E). We found no correlation between amount of exon skipping and RNA abundance, RPF levels or presence of FMRP CLIP tags at the gene level (Supplemental Figure S5 A, B and C). We next assessed the H3K36me3 levels at the skipped exons in the SE category and found a significant decrease in the H3K36me3 marks at the 5’ splice site (+/- 50nt) of these exons that are skipped in the *Fmr1* KO hippocampus compared to the wild type (Figure 7F, p-value=0.02 using the Wilcox test). Interestingly, when we increased the analysis over a region of +/-150 nt (∼2 nucleosomes) we no longer found a decrease in H3K36me3 at the skipped exons in the FMRP KO animal (Supplemental Figure S5D). Although, at the island level which encompasses mainly regions >1kb we see mainly an increase in H3K36me3 levels, at the 5’ splice site of genes (+/- 50 nt) susceptible to alternative splicing in the FMRP KO, we see a decrease in H3K36me3 levels. Further studies are needed to understand how H3K36me3 modifications and/or other factors might play a role in regulating differential alternative splicing in the FMRP KO background. Thus, as with other autism spectrum disorders, dysregulated alternative splicing is prevalent in a mouse model of FXS and might be linked to changes in H3K36me3 levels at the splice site.

**Figure 7.**
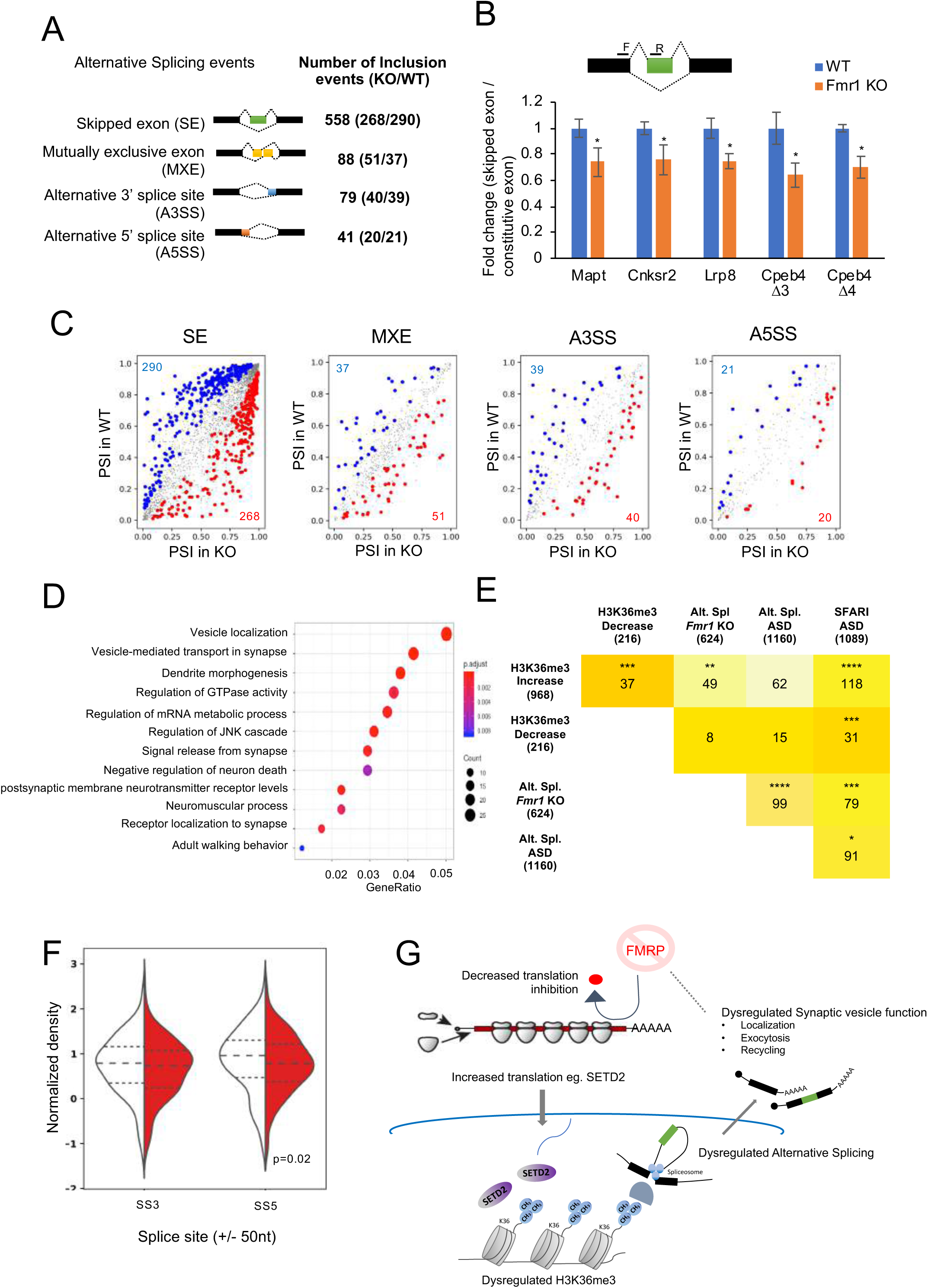
Global analysis of FMRP mediated alternative splicing (AS). (A) Summary table of total splicing events in *Fmr1* KO and WT based on RNA-seq from hippocampal slices. Splicing events detected by rMATS at a FDR < 5% and a difference in the exon inclusion levels between the genotypes (deltaPSI) ≥ 5% are depicted. (B) Alternatively spliced events validated using RT-qPCR are shown for several RNAs (hippocampus tissue, WT, n=6; *Fmr1* KO, n=6). The illustration depicts an example of exon skipping (green box) in the *Fmr1* KO. Primer positions are depicted with black bars. (C) Alternative splicing events detected in *Fmr1* KO and WT hippocampus. Inclusion events that were significantly (p-value<0.05) increased (red), decreased (blue), or unchanged (grey) are indicated. The number of events in the up or down category are shown in red and blue, respectively. PSI, (Percent Splice In / Exon inclusion levels). (D) GO term enrichment for all alternative splicing events are shown (p.adjust value <0.05). The total number of genes identified in the RNA-seq from (Figure 1) were used as background. (E) Table depicting the overlap of the alternatively spliced (Alt. Spl./AS) genes in this study with the SFARI Autism Spectrum Disorder database, the Alt. Spl. genes identified in samples from autism patients (Parikshak et al., 2016), and genes with increased or decreased H3K36me3 islands from Figure 6. The intensity of the colour represents the increasing number of overlapping genes between the gene sets. Asterisks indicate statistical significance (*,p-value<0.05; ***, p-value<0.001; ****,p-value< 0.0001, hypergeometric test). (F) Violin plot for the H3K36me3 ChIP signal at +/-50nt of the 5’ (SS5) and 3’ (SS3) splice sites of the alternatively skipped exons in WT (white) and *Fmr1* KO (red) hippocampus tissue (p-value<0.05, K-S test for significance). (G) Model for FMRP mediated alterations in H3K36me3 marks on the chromatin and alternative splicing of transcripts in the hippocampus. (also see Table S3)

## Discussion

We used several approaches to investigate FMRP regulation of mRNA expression in the mouse brain, focusing first on steady state ribosome profiling in hippocampal slices. *Fmr1* deficiency resulted in ribosome footprint levels that mostly tracked altered levels of RNA, which is consistent with other studies (Liu et al 2018; 2019; Das Sharma et al 2019). Because ribosome profiling at steady state cannot distinguish between static and translocating ribosomes, we modified our profiling procedure (Ingolia et al 2011) to identify the dynamic range of ribosome translocation rates genome-wide in hippocampal slices. These studies in turn resulted in four main novel findings. First, they showed that mRNAs cluster into defined functional biology groups based on the transit rates of their associated ribosomes. The ribosomes on most mRNAs (>5000), transit rapidly whereas on some mRNAs (∼3000), ribosome transit is slow or nearly not at all (∼350). There is some CDS length dependency as well. Second, using a combination of ribosome runoff and sucrose gradient centrifugation, we identified ∼50 mRNAs associated with FMRP-stalled ribosomes, which strongly imply translational regulation. One of these mRNAs encodes a histone lysine methyltransferase, SETD2, which is elevated in *Fmr1* KO hippocampus. Third, the SETD2-dependent histone mark, H3K36me3, is rearranged in both intragenic and intergenic regions upon *Fmr1* deficiency; primarily increases in H3K36me3 are observed in *Fmr1* KO. Fourth, because H3K36me3 has been correlated with alternative RNA processing in mammals (Kim et al 2011), we analyzed our RNA-seq data and found widespread aberrant splicing in the *Fmr1* KO hippocampus, most prominently exon skipping/inclusion. These altered processing events often correlated with changes in the histone mark, occurred on genes that regulate pre-synaptic functions such as synaptic vesicle recycling, and were strongly linked to autism spectrum disorders (Figure 7E).

Earlier studies have inferred that ribosome stalling in the brain is linked to RNA transport granules in dendrites (Krichevsky and Kosik 2001; Mallardo et al 2003; Kanai et al 2004; Khandjian et al 2004). It is possible that at least some of the ribosome stalling we observe may be the result from such compartmentalization in neurons, or perhaps is a more general feature of the brain. In contrast, rapidly dividing ES cells do not appear to have stalled ribosomes, at least not of the magnitude we observe here (Ingolia et al 2011). Irrespective of whether ribosome stalling is a dendritic or neuronal phenomenon, how it occurs mechanistically on specific mRNAs is unclear, but it could be related to secondary structure or association with RNA binding or other proteins. One clue, however, could be the accumulating evidence that strongly implicates phosphorylation of eEF2 as a coordinator of translation and neural activity, and thus is likely to be one mediator of ribosome translocation rates (Scheetz et al 2000; Sutton et al 2007; Park et al 2008; McCamphill et al 2015; Heise et al 2017). Even so, the involvement of phospho-eEF2, a general translation factor, would unlikely be responsible for mRNA-specific stalling. We also noted that some mRNAs, such as those in sub-cluster 2 (Figure 3D, E) encode proteins involved in the extracellular matrix (Figure S2C). A number of these proteins contain proline-rich stretches, whose geometry in the polypeptide exit tunnel can lead to ribosome stalling (Schuller et al 2017; Hutter et al 2017).

We have identified 48 mRNAs whose ribosomes are stalled specifically by FMRP. These RNAs include those with (40 mRNAs) and without FMRP CLIP (8 mRNAs) sites in coding regions (Darnell et al 2011, Driesche et al., 2019), so it would appear that a simple FMRP “roadblock” model would not explain ribosome stalling. Alternatively, Ishizuka et al (2002) showed that *Drosophila* FMRP binds ribosomal proteins, and in a cryo-EM study, Chen et al (2014) found that *Drosophila* FMRP reconstituted with mammalian ribosomes binds a region that would preclude tRNA and elongation factors from engaging in translation. However attractive this model is for FMRP-mediated ribosome stalling, the fact that mRNA or translation factors were not included in the reconstitution assays brings into question whether it reflects the physiological basis for ribosome stalling.

Our studies revealed that FMRP stalls ribosomes on a number of mRNAs that encode epigenetic regulators and that the mis-regulation of one of them in *Fmr1*-deficient hippocampus, SETD2, leads to widespread increase in H3K36me3 chromatin marks (islands). These H3K36me3 islands are both enriched for and depleted from autism risk genes (Figure 6E) as well as those with synaptic functions (Figure 6F). This suggests that SETD2, the main enzyme that catalyzes this chromatin mark, might also be an autism risk gene, which has been previously reported (Lumish et al 2015). Mutations in several chromatin remodeling genes have been linked to changes in neural function, autism, and other intellectual disabilities (van Bokhoven 2011; Goodman and Bonni 2019). In most cases, however, mutations in epigenetic modifiers result in up or down regulation of transcription. Indeed, previously identified epigenetic changes in FXS model mice by Korb et al (2017), particularly H3K4me3, H4K8ac, and H4K16ac, were responsible for widespread changes in transcription in cortical neurons and the cerebellum. Importantly, these investigators could rescue several FXS pathophysiology’s, many of which resemble those in autism, by targeting the epigenetic reader protein Brd4. H3K36me3 does not mark promoter regions, but instead is found in gene bodies and increases as RNA polymerase II proceeds 3’ during catalysis (Neri et al 2017). In mammals, one of the functions H3K36me3 is correlated with, if not causative for, is alternative pre-mRNA processing (Kim et al 2011). In the *Fmr1*-deficient hippocampus, we detected extensive changes in RNA processing, mostly exon skipping. Surprisingly, most of the skipped exons occurred in genes that have pre-synaptic functions such as synaptic vesicle recycling, transport and exocytosis which could at least partially explain alterations in synaptic vesicle dynamics and neurotransmitter release in FXS model mice (Deng et al 2013; Ferron et al 2014; Broek et al 2016). Furthermore, a number of skipped exons in *Fmr1* KO have been linked to neurological disorders such as autism (skipped microexon 4 in *Cpeb4* (Irimia et al., 2014; Parras et al., 2018; Quesnel-Vallières et al., 2016)), Alzheimer’s disease (skipped exon 7 in *APP* (Fragkouli et al., 2017; Rockenstein et al., 1995)), Parkinson’s disease (skipped exon 3 in *Mapt* (Lai et al., 2017)), and intellectual disabilities (skipped exon12 in *Cnksr2* (Houge et al., 2012)).

Alternative pre-mRNA processing is a common feature in autism (Quesnel-Vallieres et al 2019), so it is perhaps not surprising that this would occur in Fragile X. However, because FMRP is an RNA binding protein that regulates translation, a reasonable assumption would be that mRNAs encoding splicing factors are improperly expressed in the disorder, and that this leads directly to mis-splicing events. However, our data indicate that this relationship might not be linear. Instead, we find that FMRP controls ribosome dynamics on specific mRNAs, which leads to elevated translation (e.g., SETD2), which leads to chromatin modifications (e.g., H3K36me3), which in turn might lead to alternative splicing of neuronal mRNAs (Figure 7G). These observations do not preclude the possible, if not likely, direct involvement of FMRP in some of these events. Even so, inferences that Fragile X is a disorder of improper translation, while not incorrect, is vastly oversimplified. However, the complexity of gene expression changes in Fragile X might offer opportunities for therapeutic intervention at multiple steps.

## Acknowledgements

We thank Lori Lorenz, Craig Peterson, Huan Shu and Danesh Moazed for valuable discussions. JDR, KMH, BL, and SS conceived various aspects of the project. We thank Maria Ivshina and Suna Jung for providing reagents. This work was supported by grants from the NIH (U54HD82013 to JDR, U54HD082008 to KMH), the Simons Foundation for Autism Research Initiative (to JDR and KMH), and the Charles H. Hood Foundation (to JDR). SS was supported by a postdoctoral fellowship from the FRAXA Foundation.

## Author Contributions

BL, GM, SS performed the experiments. RW provided essential help with bioinformatics. JDR wrote the manuscript with input from all authors.

## LEAD CONTACT AND MATERIALS AVAILABILITY

Further information and requests for resources and reagents should be directed to and will be fulfilled by the Lead Contact, Joel Richter. This study did not generate new unique reagents.

## STAR METHODS

### EXPERIMENTAL MODEL AND SUBJECT DETAILS

All experiments were approved by the Institutional Animal Care and Use Committee at University of Texas Southwestern and University of Massachusetts Medical School and conducted in accordance with the National Institutes of Health Principles of Laboratory Animal Care. Fmr1 KO mice were obtained from Dr. Stephen T. Warren and backcrossed for more than 10 generations on a C57BL/6 background. Food and water were provided *ad libitum*, and mice were kept on a 12 h light/dark cycle (7am-7pm light period). Pups were kept with their dams until weaning at postnatal day 21. After weaning, mice were group housed (maximum of 6 per cage) by sex. Cages were maintained in ventilated racks in temperature (20-21°C) controlled rooms. P28-35 wild-type or *Fmr1* KO male mice littermates were used in this study.

#### Mouse acute brain slice preparation

Transverse hippocampal or hippocampal-cortical brain slices were acutely prepared from P28-35 C57BL/6N wild-type or *Fmr1* KO male mice littermates as previously described (Guo et al., 2016). Briefly, mice were anesthetized with ketamine (125 mg/kg)/xylazine (25 mg/kg) and transcardially perfused with chilled (4°C) sucrose dissection buffer containing the following (in mM): 2.6 KCl, 1.25 NaH_2_PO_4_, 26 NaHCO_3_, 0.5 CaCl_2_, 5 MgCl_2_, 212 mM sucrose, and 10 dextrose aerated with 95% O_2_/5% CO_2_. Transverse hippocampal slices (400µm) were obtained on a Leica VT1200S slicer. For ribosomal profiling experiments (Figure 1), CA3 was cut off to enrich for CA1 and for each biological replicate, 8-10 slices from 6-8 mice/genotype were pooled. Slices were recovered and maintained at 30°C for 3-4 hours in artificial cerebrospinal fluid (ACSF) which contained the following (in mM): 119 NaCl, 2.5 KCl, 2 CaCl_2_, 1 MgCl_2_, 26 NaHCO_3_, 1 NaH_2_PO_4_ and 11 D-glucose aerated with 95%/O_2_/5% CO_2_ to pH 7.4. Slices were then treated with cycloheximide (100 µg/ml) in ACSF, 4°C, snap-frozen on dry ice/EtOH bath, and stored at -80°C. For homoharringtonine (HHT) experiments (Figures 3-5), cortical-hippocampal slices were recovered 3-4 hours in ACSF and then treated with HHT (20 µM) (Tocris) or 0.05% DMSO (vehicle) for different times (0, 5, 10, 30, 60 min). At the end of the incubation time, slices were snap/frozen on dry ice/EtOH bath and stored at -80 C. Ten slices from 2-3 mice/genotype were pooled for each biological replicate.

##### Cell culture

Immortalized Mouse Embryonic Fibroblasts (MEFs) were derived from C57BL/6N wild-type or *Fmr1* KO as described earlier (Groisman et al., 2012). Immortalized MEF cells were cultured in DMEM (Dulbecco’s Modified Eagle’s Medium, Gibco) with 10% fetal bovine serum using the 3T3 culture protocol and cells derived from Passage number 31 onwards were used for western blot analysis.

### METHOD DETAILS

#### Sucrose gradients of hippocampal slices for steady-state ribosome profiling

Frozen, isolated CA1 hippocampal slices were thawed in ice-cold homogenization buffer (20mM Tris-HCl pH7.4, 5mM MgCl_2_, 100mM KCl, 1mM DTT, 100μg/ml CHX (cycloheximide), 25U/ml Turbo DNaseI (Ambion, #AM2238), 1X EDTA-free protease inhibitor (Roche), avoiding detergent in nuclease-free water) on ice for 5min. Wide orifice tips were used to transfer slices to a pre-chilled detergent-free Dounce homogenizer. Tissues were slowly homogenized by hand (20 strokes of loose pestle A, and 20 strokes of tight pestle B). Homogenates were carefully transferred to clean 1.5ml tubes with clean glass Pasteur pipets and bulbs. 1% NP-40 was added to the homogenates and incubated on ice for 10min. Homogenates were clarified by centrifugation at 2,000g 4 °C for 10min. The supernatants were collected and clarified again by centrifugation at 20,000g 4 °C for 10min. The supernatants were collected, and the amounts of nucleic acid were measured by Nanodrop (A_260_ units). For each sample, cytoplasmic RNA for RNA-seq was purified from one-fourth of the lysate with TRIzol LS reagent (Invitrogen, #10296028). The other three-fourths of the lysate was digested with 100ng RNase A (Sigma, # R4875) and 60U RNase T1 (Thermo Fisher Scientific, #EN0542) per A_260_ at 25°C for 30 min and stopped by chilling on ice and adding 50U SUPERase In RNase inhibitor (Ambion, #AM2694). Digested lysates were applied to 10%-50% (w/v) sucrose gradients prepared in 1X polysome buffer (20mM Tris-HCl pH7.4, 5mM MgCl_2_, 100mM KCl, 1mM DTT, 100μg/ml CHX in nuclease-free water). After the ultracentrifugation in a SW41Ti rotor (Beckman Coulter) at 35,000 rpm (avg 151,263g) 4°C for 2.5 hours, gradients were fractionated at 1.5 ml/min and 12 sec collection intervals through a fractionation system (Brandel) that continually monitored A_260_ values. Monosome fractions were identified, pooled, and extracted with TRIzol LS.

#### Sucrose gradients of cortical-hippocampal slices for run-off ribosome profiling

Frozen cortical-hippocampal slices were thawed in ice-cold lysis buffer (20mM Tris-HCl pH7.4, 5mM MgCl_2_, 100mM KCl, 1mM DTT, 100μg/ml CHX, 25U/ml Turbo DNaseI, 1X EDTA-free protease inhibitor, 1% NP-40, in nuclease-free water) on ice for 5min. Tissues were dissociated by pipetting and further trituration through 25G needle for 10 times. Lysates were incubated on ice for 10min and then clarified by centrifugation at 2,000g 4 °C for 10min. The supernatants were collected and clarified again by centrifugation at 20,000g 4 °C for 10min. The supernatants were collected, and the amounts of nucleic acid were measured with Qubit HS RNA assays. Lysates containing ∼40µg RNA were digested with 600ng RNase A (Ambion, #AM2270) + 75U RNase T1 (Thermo Fisher Scientific, #EN0542) /µg RNA in 0.3ml at 25°C for 30min and stopped by chilling on ice and adding 50U SUPERase In RNase inhibitor. Digested lysates were applied to 10%-50% (w/v) sucrose gradients similarly as above.

#### Sucrose gradients of cortical-hippocampal slices for polysome profiling

Frozen cortical-hippocampal slices were thawed in ice-cold homogenization buffer on ice for 5min. Wide orifice tips were used to transfer slices to a pre-chilled detergent-free Dounce homogenizer. Tissues were gently homogenized by hand (3 strokes of loose pestle A, and 7 strokes of tight pestle B). Homogenates were carefully transferred to clean 1.5ml tubes with clean glass Pasteur pipets and bulbs. Homogenates were clarified by centrifugation at 2,000g 4 °C for 10min. The supernatants were collected and clarified again by centrifugation at 10,000g 4 °C for 10min. The supernatants were collected, and the amounts of nucleic acid were measured with Nanodrop (A_260_ units). 1% NP-40 was added to the homogenates and incubated on ice for 10min. ∼1A_260_ unit of lysate was saved for the input RNA. ∼5 A_260_ units of lysate were applied to 15%-45% (w/w) sucrose gradients prepared in 1X polysome buffer and with the lower block mark and long cap. After the ultracentrifugation in a SW41Ti rotor (Beckman Coulter) at 35,000 rpm (avg 151,263g) 4°C for 2 hours, gradients were fractionated at 1.5 ml/min and 20 sec collection intervals through a fractionation system (Brandel) that continually monitored A_260_ values. Some gradients were treated with RNase A prior to centrifugation.

#### Ribosome profiling

Ribosome profiling libraries were prepared following the published protocols (Heyer et al., 2015). Briefly, rRNA was depleted from the purified monosomal RNA with RiboZero (Illumina, #MRZG12324). Remaining RNA samples were separated on a 15% TBU gel (National Diagnostics, #EC-833) and the ribosome footprints were size-selected between the 26 and 34nt markers. RNA was eluted from the crushed gel pieces in RNA elution buffer (300mM NaOAc pH5.5, 1mM EDTA, 0.25% SDS) at room temperature overnight, filtered with Spin-X Centrifuge Tube Filters (Corning, #8162) and precipitated with equal volume of isopropanol. Recovered RNA was dephosphorylated with T4 Polynucleotide Kinase (NEB, #M0201S) and ligated with preadenylated adaptor in miRCat®-33 Conversion Oligos Pack (IDT) using T4RNL2Tr.K227Q ligase (NEB, #M0351L). Reverse transcription (RT) was performed with primers containing 5nt-barcodes and 8nt-unique molecular identifiers (UMIs) and SuperScript III (Invitrogen, #18080-044) in 1X first-strand buffer without MgCl_2_ (50 mM Tris-HCl, pH 8.3, 75 mM KCl). RT products were separated on a 10% TBU gel and the 130-140nt region was selected. cDNA was eluted in DNA elution buffer (10mM Tris pH 8.0, 300mM NaCl, 1mM EDTA) at room temperature overnight, filtered, and precipitated with isopropanol. Purified cDNA was circularized with CircLigase (Epicentre, #CL4115K). Except for the RNase titration samples, cDNA derived from remaining rRNA was hybridized to biotin-labelled antisense probes (IDT) and further depleted with Dynabeads MyOne Streptavidin C1 (Invitrogen, #65001). Optimal PCR cycle number was determined empirically by test PCR reactions with titrated cycle numbers. Final PCR amplification was performed with KAPA Library Amplification Kit (Kapa Biosystems, #KK2611) and 180-190bp products were size-selected on an 8% TBE gel. DNA was eluted in DNA elution buffer, filtered, and precipitated with isopropanol. The final library DNA was purified with AMPure XP beads (Beckman Coulter, #A63880). Oligos used for the library preparation are listed in Table S2.

In parallel, input RNA samples were processed similarly as the ribosome footprints except the following steps. After rRNA depletion, input RNA mixed with an equal volume of 2x alkaline fragmentation solution (2 mM EDTA, 10 mM Na_2_CO_3_, 90 mM NaHCO3, pH ∼ 9.3) and heated at 95°C for 15min to achieve an average fragment length of ∼140nt. Fragmented RNA samples were separated on a 10% TBU gel and size-selected between the 100 and 150nt markers. RT products were separated on a 10% TBU gel and the 200-250nt region was selected. Antisense probe depletion was omitted for input RNA samples. 260-300bp final PCR products were size-selected on an 8% TBE gel.

The size distributions of final libraries were measured by Fragment Analyzer (Advanced Analytical, performed by Molecular Biology Core Labs at UMMS). The concentrations were quantified with KAPA Library Quantification Kit (Kapa Biosystems, #KK4835). Libraries were pooled with equal molar ratios, denatured, diluted, and sequenced with NextSeq 500/550 High Output Kit v2 (Illumina, 75bp single-end runs, #FC-404-2005) on a Nextseq500 sequencer (Illumina).

#### RNA-seq for polysome fractions

Based on the FMRP signals in the immunoblotting of gradient fractions, medium and heavy fractions were identified, pooled, and extracted with TRIzol LS together with saved input lysates. RNA samples were quantified with Qubit HS RNA assay kit and the integrity was examined with Fragment Analyzer. 200ng RNA was used for BIOO Rapid directional qRNA-seq library preparation following the manufacturer’s instructions. 12 cycles were used for the final PCR amplification. The libraries were quantified with KAPA Library Quantification Kit and the quality was examined with Fragment Analyzer. Libraries were pooled with equal molar ratios, denatured, diluted, and sequenced with NextSeq 500/550 High Output Kit v2 (Illumina, 80bp pair-end runs for RNA-seq, #FC-404-2002) on a Nextseq500 sequencer (Illumina).

#### Western blotting

Hippocampi were homogenized at 4°C in RIPA buffer. Protein complexes were released by sonication at 4°C and the extract was centrifuged at 13,200 rpm for 10 min at 4°C and the supernatant collected. Protein concentration was determined by BCA reagent. Proteins (10 μg) were diluted in SDS-bromophenol blue reducing buffer with 40 mM DTT and analyzed using western blotting with the following antibodies: SETD2 (ABclonal, A11271) and Lamin AC (Thermo Fisher, 14-9688-80). After incubation in primary antibody, immunoblots were incubated with HRP-conjugated secondary antibodies (Jackson Immunoresearch) and developed by Clarity ECL substrate (Biorad). For western blots using MEFs, cells were harvested and lysed in Triton Extraction Buffer and nuclear protein extracts were obtained as above and detected by SETD2 and Lamin B1 (Abcam, ab16048) antibodies. For sucrose gradient fractions, 60µl samples were mixed with 20µl 4X SDS loading dye (240mM Tris-HCl pH 6.8, 5% beta-mercaptoethanol, 8% SDS, 40% glycerol, 0.04% bromophenol blue) and boiled at 95°C for 10min. Samples were briefly heated at 95°C again for 30sec immediately before loading on 10% SDS-PAGE gel (35 µl/sample). Separated proteins were transferred to PVDF membranes at constant 90 mA 4°C for 16 hours. Membranes were blocked with 5% non-fat milk in 1X TBST at room temperature for 1hour and incubated with primary antibody at 4°C overnight. FMRP (Abcam, 1: 2000), Tubulin (Sigma, 1:5000), Rpl4 (Proteintech, 1:5000), UPF1 (Abcam, 1: 5000), MAP2 (Millipore, 1:2000), eEF2 (Cell signaling, 1:2000), GAPDH (Cell signaling, 1:2000), MRPS18B (Proteintech, 1:2000) and Rps6 (Cell signaling, 1:4000) were diluted in 1X TBST with 5% non-fat milk. Membranes were washed three times for 10min with 1XTBST and incubated with anti-rabbit or anti-mouse secondary antibodies (Jackson, 1:10000) at room temperature for 1hour. Membranes were washed three times for 10min with 1XTBST, developed with ECL-Plus (Piece), and scanned with GE Amersham Imager

#### Chromatin immunoprecipitation Sequencing (ChIP-Seq)

ChIP was performed as previously described (Cotney and Noonan, 2015). Briefly, hippocampal tissue was isolated from 4 adult mice (P35) per genotype and minced into tissue < 0.5mm^3^ in 250ul of ice-cold PBS with protease inhibitors. Tissue was cross-linked with 1% formaldehyde and rotated at room temperature for 15 min at 50 rpm and quenched with 150mm glycine for 10 min in a total volume of 1ml PBS. The supernatant was discarded after centrifugation at 2000g for 10 min at 4^0^C and the pellet was resuspended in 300ul of chilled cell lysis buffer for 20 min on ice. Swollen pellets were homogenized using a glass Dounce homogenizer (2ml) with 40 strokes of a tight pestle on ice. Nuclei were harvested after centrifugation at 2000g for 5min at 4^0^C and resuspended in 300ul ice-cold nuclear lysis buffer for 20min on ice. SDS was added to make a final concentration of 0.5%. Samples were sonicated on a Bioruptor® sonicator at high power settings for 9 cycles (sonication: 30 sec on, 90 sec off) of 15min each at 4°C at high power. Supernatants were collected after centrifugation at 16,000g for 10 min at 4^0^C. The samples were diluted to bring the SDS concentration <0.1%; 10% of each sample was reserved as input. The rest of the samples were divided into two and incubated with Protein G dynabeads coupled overnight at 4^0^C with either H3K36me3 (Abcam ab9050, 5μg per ChIP) or IgG (Sigma 12-371, 5 μg per ChIP). After IP, beads were washed, and chromatin eluted in elution buffer for 20 min shaking at 65**°**C. IP and Input samples were de-crosslinked overnight at 65**°**C. RNAse digestion for 1 hr. at 37**°**C and proteinase K treatment for 30 min at 55**°**C was performed. DNA was then purified with the QIAGEN PCR purification kit in 50ul elution buffer. For library preparation, purified input and IP DNA was end repaired using T4 DNA polymerase, Klenow polymerase and T4 Polynucleotide kinase from NEB at 20**°**C for 30 min. DNA was extracted using 35ul of the Agencourt Ampure XP beads and ‘A’ bases were added to the 3’ end using Klenow exonuclease (3’ to 5’ exo minus) from NEB for 30min at 37**°**C. DNA was purified using 60ul of the Agencourt Ampure XP beads and Illumina adapter sequences were ligated to the DNA fragments using the Quick Ligase (NEB) for 15min at 20**°**C. The library was size-selected using 50ul of Agencourt Ampure XP beads. Using the multiplexing barcoded primers, the library was PCR amplified and purified using 50ul of the Agencourt Ampure XP beads and analyzed using a Fragment Analyzer (Advanced Analytical, performed by Molecular Biology Core Labs at UMMS). Libraries were pooled with equal molar ratios, denatured, diluted, and sequenced with NextSeq 500/550 High Output Kit v2.5 (Illumina, 75bp paired-end runs,) on a Nextseq500 sequencer (Illumina).

### QUANTIFICATION AND STATISTICAL ANALYSIS

All grouped data are presented as mean ± s.e.m. All tests used to compare the samples are mentioned in the respective figure legends and corresponding text. When exact p values are not indicated, they are represented as follows: *, p < 0.05; **, p < 0.01; ***, p < 0.001; ****, p-value <0.0001; n.s., p > 0.05.

#### Ribosome profiling analysis

Individual samples were separated from the raw fastq files based on the barcode sequences. Adaptor sequences (TGGAATTCTCGGGTGCCAAGGAGATCGGAAGAGCGGTTCAGCAGGAATGCCGAGACCG) were removed with cutadapt (1.7.1). Trimmed fastq files were uploaded to the Dolphin platform (https://www.umassmed.edu/biocore/introducing-dolphin/) at the UMMS Bioinformatics Core for the mapping steps. Trimmed reads were quality filtered with Trimmomatic (0.32) and mapped to the mouse rRNA and then tRNA references with Bowtie2 (2.1.0). Unmapped reads were next mapped to the mm10 mouse genome with Tophat2 (2.0.9). Reads mapped to >1 loci of the genome were classified as “multimapped” reads and discarded. PCR duplicates were marked based on the UMI sequences with custom scripts and only uniquely mapped reads without duplicates were retained with Samtools (0.0.19) for the downstream analyses. RPF length distribution, P-site offsets, and frame preference were calculated with plastid (0.4.8). Counts at each nucleotide position were extracted using P-sites of RPFs and 5’end of mRNA reads with +11 offset, normalized to the library size, averaged across replicates, and plotted along mRNA positions with custom scripts.

#### Steady-state differential translation analysis

Cleaned bam files were converted to fastq files with bedtools. For both ribosome profiling and RNA-seq, gene expression was quantified with RSEM (1.2.11) using the cleaned fastq files and Refseq (V69) mouse CDS without the first and last 30nt to avoid the translation initiation and termination peaks. Genes were filtered with minimum 10 reads across all replicates and then the read counts were batch-corrected with the Combat function in sva (3.24.4) using a full model matrix. Batch-corrected counts were normalized with trimmed mean of M values (TMM) method and used to identify differential expressed genes (DEGs) with anota2seq (1.0.0). Instead of the default setting, the priority of TE groups were set to be higher than that of mRNA only groups. A permutation test was performed to estimate the false discovery rate (FDR) with nominal p-value < 0.01, which was 0.097. GO analysis was performed and plotted with clusterProfiler (3.10.1) using all genes past filtering in the dataset as the background.

#### Hierarchical clustering of ribosome run-off and gene ontology (GO) enrichment of each cluster

RPFs were mapped as described above and the most abundant isoform estimated by RSEM for each gene was selected as the representative transcript. For the metagene analysis, transcripts with CDS longer than 3000nt (N=1401) were selected to ensure incomplete ribosome run-off after 10min HTT treatment. The read densities at each nucleotide position were normalized to the average density of the last 500nt of CDS and averaged using the P sites of RPFs. For visualization purposes, the curves were smoothed within a 90nt window. An arbitrary 0.8 threshold was used to estimate the relative run-off distances at 5 and 10min. A linear regression analysis between the HHT treatment time and ribosome run-off distances was performed to estimate the global elongation rate. To visualize the run-off patterns for individual transcripts, RPFs were normalized to the abundance of mitochondrial RPFs and plotted along the mRNA nucleotide positions. The same y-scale was used across the time-course, so the initiation peaks were truncated to reveal the patterns over CDS. Stalled ribosomes were quantified by RSEM by mapping RPFs to Refseq (V69) mouse CDS without the first and last 300nt to avoid the translation initiation and termination peaks after prolonged HTT treatment. Counts of RPFs were normalized to the length of CDS minus 600nt and the abundance of mitochondrial RPFs in that sample to calculate the RPK (Reads Per Kilobase of transcript) value for each transcript.

To categorize and group genes with distinct ribosome run-off patterns, the RPF of each gene at each time point was first normalized to time point 0. The Euclidean distance matrix was then calculated, followed by hierarchical clustering using Ward’s agglomeration method (Ward, 1963). The clustering process was performed using the hclust (Galili, 2015), and the clustering heatmap was displayed using pheatmap (Kolde and Kolde, 2015) in R. To test the reliability of clustering, analysis of group similarities (ANOSIM) test (Clarke, 1993) was performed using vegan (Dixon, 2003) in R with 1000 permutation times on both overall clusters and sub-clusters. The global pattern of each cluster was summarized using the corresponding median and standard deviation in each timepoint.

GO enrichment analysis for each sub-cluster were performed through clusterProfiler (Yu et al., 2012) using genes covered by run-off ribosome profiling as background. The statistical significance was adjusted using FDR. To remove redundancy in reporting, each reported GO term was required to have at least 25% of genes that were not associated to another term with a more significant p value.

#### Polysome fraction RNA-seq data analysis

Fastq files were uploaded to the Dolphin platform at the UMMS Bioinformatics Core for mapping and quantification. 9nt molecular labels were trimmed from both 5’ends of the pair-end reads and quality-filtered with Trimmomatic (0.32). Reads mapped to mouse rRNA by Bowtie2 (2.1.0) were filtered out. Cleaned reads were next mapped to the Refseq (V69) mouse transcriptome and quantified by RSEM (1.2.11).

Estimated counts on each gene were used for the differential gene expression analysis by DESeq2 (1.16.1). After the normalization by median of ratios method, only the genes with minimal 5 counts average across all samples were kept for the DEG analysis. Data were fit to a model of treatment+fraction+treatment:fraction+treatment:genotype:fraction and the treatmenthht.fractionheavy result was used to identify mRNA enriched or depleted in the heavy fractions after HTT treatment. P<0.05 and 2-fold change were used as the cut-offs.

For the medium fractions after HHT treatment, *Fmr1* KO samples was directly compared to the WT samples and p<0.05 was used as the cut-off for identifying DEGs. Steady state ribosome profiling data of the decreased DEGs were compared to all the other genes to independently confirm whether these mRNAs indeed had fewer stalled ribosomes in *Fmr1* KO.

#### ChIP-Seq analysis

For ChIP-sequencing analysis, alignments were performed with Bowtie2 (2.1.0) using the mm10 genome, duplicates were removed with Picard and TDF files for IGV viewing were generated using a ChIP-seq pipeline from DolphinNext (Yukselen et al., 2019). The broad peaks for H3K36me3 ChIP-Seq were called using the SICER v1.1 package (Xu et al., 2014). H3K36me3 enriched islands were identified using the parameters set to a window size of 200bp, gap size 600bp and FDR cutoff of 1 × 10^-3^, and the default value for the redundant rate cutoff. To analyze the H3K36me3 difference level between WT and FMRP KO, the reads count on each island of each replicate was first normalized to total mapped reads and then normalized to its corresponding input sample. The H3K36me3 difference level of each island is then calculated using both replicates in FMRP KO and WT groups; a negative binomial test is performed using edgeR (Robinson et al., 2009) to consider variability between replicates and differences between FMRP KO and WT groups. The islands with significantly different H3K36me3 levels are defined using a cut-off for fold change >2 and p-value < 0.05. Gene Ontology (GO) enrichment analysis was performed using the clusterProfiler package (Yu et al., 2012) to obtain GO terms related to Biological processes in the genes with differentially enriched islands in the *Fmr1* KO vs Wild type. deepTools was used to plot heatmaps and profiles for genic distribution of H3K36me3 ChIP signal. Samtools (0.1.19) was used for sorting and converting Bam files. RPGC (per bin) = number of reads per bin / scaling factor for 1x average coverage was used for normalization where the scaling factor was determined using sequencing depth: (total number of mapped reads x fragment length) / effective genome size. IGV tools (2.3.67) was used for visualizing TDF files and all tracks shown are normalized for total read coverage.

To quantify the H3K36me3 level at the 5’ and 3’ splicing sites of the SE, the read density at a region of ±50nt at each splice site (total 100nt) and ±150nt at each splice site (total 300nt) was normalized to the read density in the entire gene body, which controls for any fluctuations in total H3K36me3 at the respective genes between both genotypes (p-value calculated using the Wilcox test for significance

#### Alternative splicing analysis

RNA-seq from hippocampal slices was used to analyze alternative splicing (AS) using the rMATS package v3.2.5 (Shen et al., 2014) with default parameters and reads were trimmed to 50bp. The Percent Spliced In (PSI) levels or the exon inclusion levels calculated by rMATS using a hierarchical framework. To calculate the difference in PSI between genotypes a likelihood-ratio test was used. AS events with an FDR < 5% and |deltaPSI| ≥ 5% as identified using rMATS were used for further analysis. We included genes that had at least one read at the differential splice junction for both genotypes. GO enrichment analysis was done by the clusterProfiler package (Yu et al., 2012) to obtain GO term enrichment for the AS events. The genes with significant skipped exons were used for validation using RT-qPCR analysis. One ug of RNA from hippocampal tissue was used to generate cDNA using the Quantitech two-step cDNA synthesis kit. Primers were designed to overlap skipped/inclusion exon junctions and qPCR was performed using the Bio-Rad SYBR reagent on a Quantstudio3 instrument.

### DATA AND CODE AVAILABILITY

Codes and scripts used for quantification analysis were written in Python or R and will be provided upon request to the Lead Contact. Data Resources Sequencing datasets generated in this study have been deposited into the Gene Expression Omnibus (GEO) database under the accession number: Super series GSE143333

---

**Figure S1.**
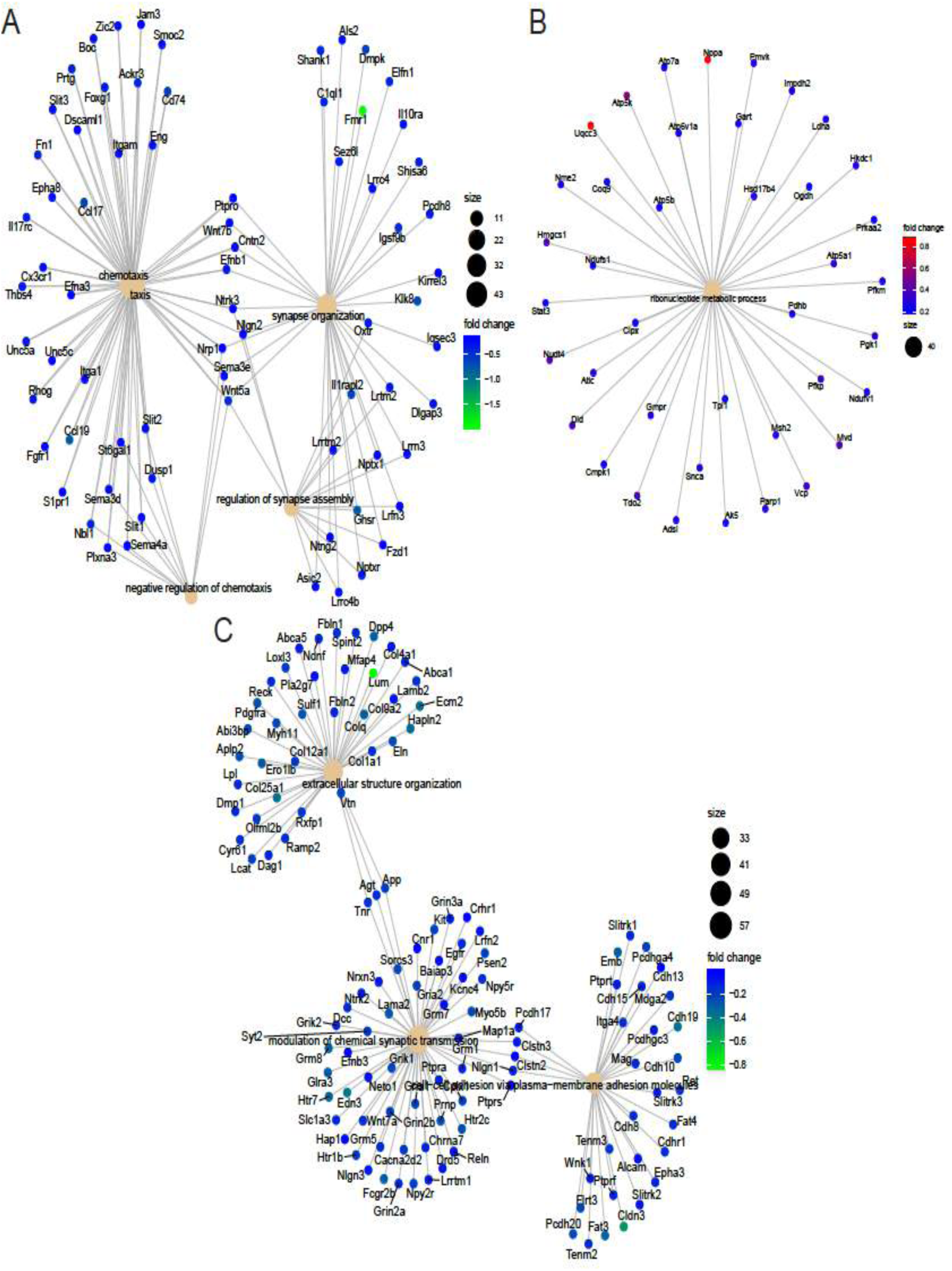
Clustering of RNAs with changes in TE and mRNA abundance to identify functional hubs (Related to Figure 1, also see TableS1) (A) Network clustering of “mRNA up” RNAs from the ribosome profiling and RNA-seq in WT and *Fmr1* KO hippocampus. Nodes indicate the functional processes linked to the RNAs identified. Fold changes are indicated. (B) Network clustering of the functional processes in the “mRNA down” RNAs (C) Network analysis of the of the functional processes in the “TE down” mRNAs

**Figure S2.**
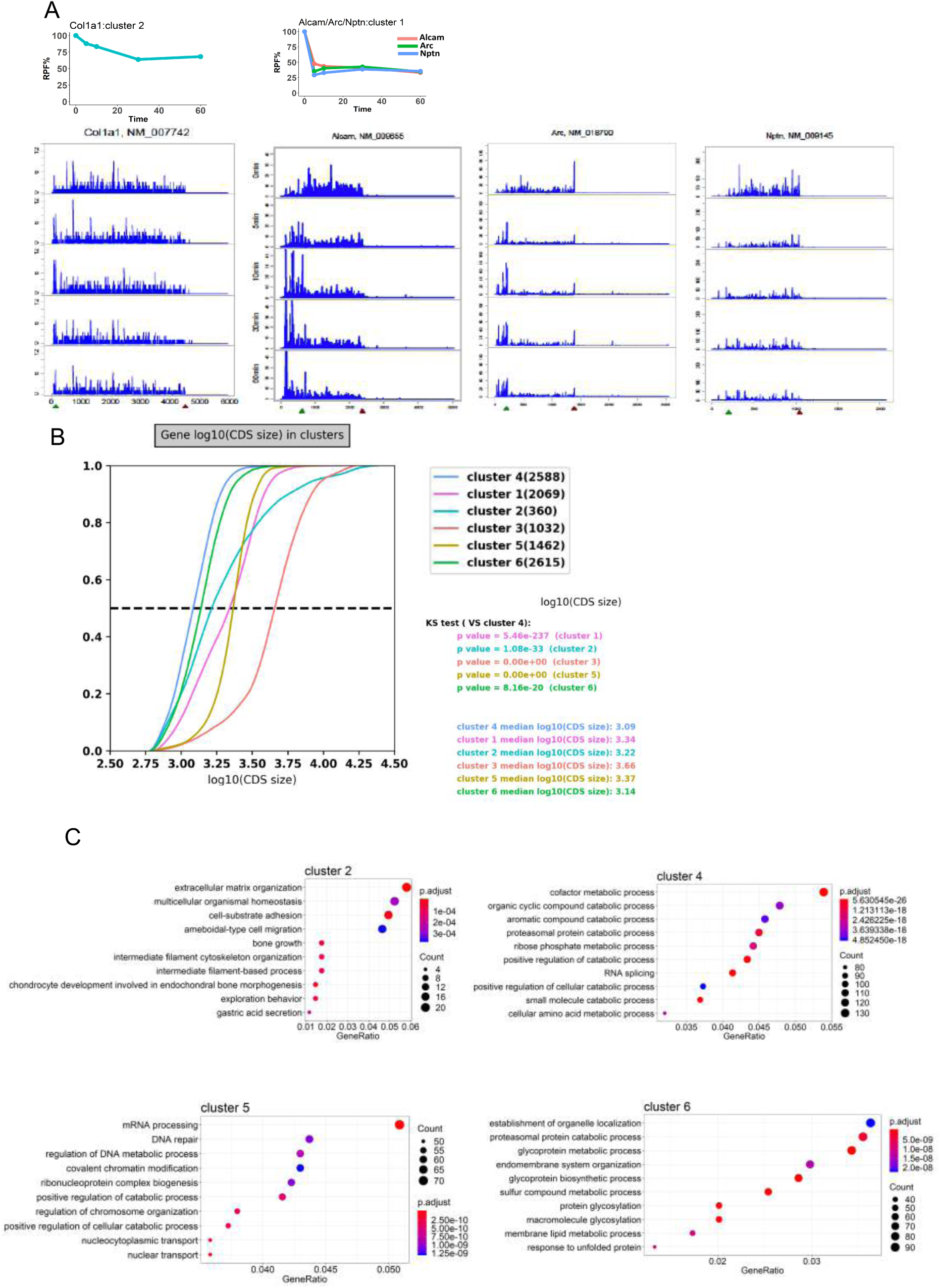
Run-off ribosome profiling of WT mouse hippocampal slices. (Related to Figure 3, also see Table S1) (A) Ribosome runoff patterns for clusters 1 and 2. The RPFs of each gene at each time point was normalized to time 0. The euclidean distance matrix was then calculated, followed by hierarchical clustering using Ward’s agglomeration method (Ward, 1963). The global pattern of each sub-cluster was summarized using the corresponding median and standard deviation in each timepoint. Representative ribosome runoff profiles are shown. (B) CDS length dependency of the ribosome runoff rates. Results of the K-S test for significance are shown (C) GO terms for sub-clusters 2,4, 5 and 6.

**Figure S3.**
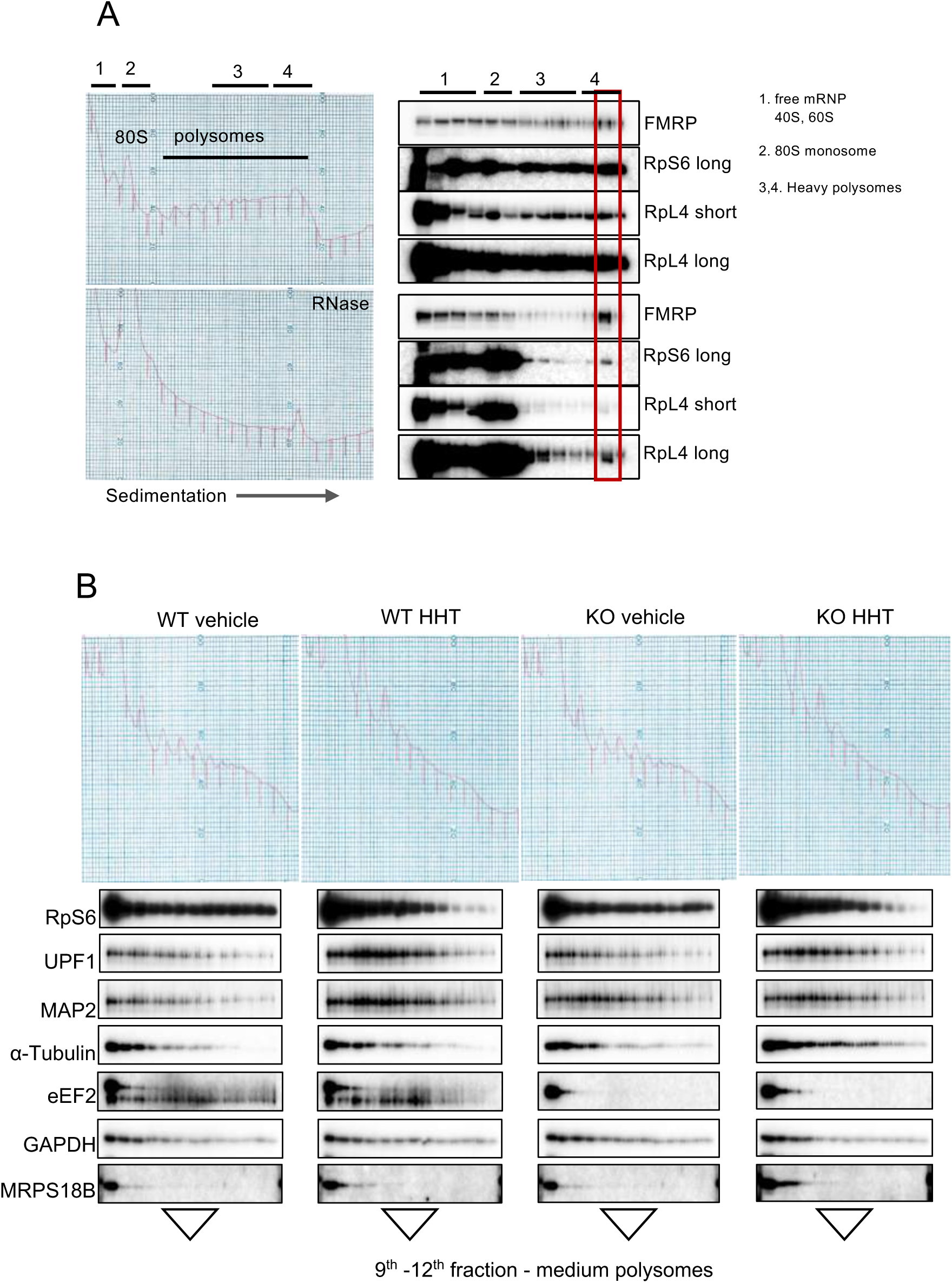
Sedimentation of proteins in polysome sucrose gradients. (Related to Figure 4, also see Table S1) (A) Hippocampal extracts, some of which were treated with RNAse A, were centrifuged through sucrose gradients, fractionated, and immunoblotted for FMRP, RPS6 and RPL4. Long and short refer to relative exposure times. (B) Hippocampal slices from WT or FMRP KO mice were treated with vehicle only or HHT for 30 min. The gradients were fractionated and immunoblotted for the indicated proteins. The red arrow denotes the eEF2 band. The medium polysome fractions are indicated.

**Figure S4.**
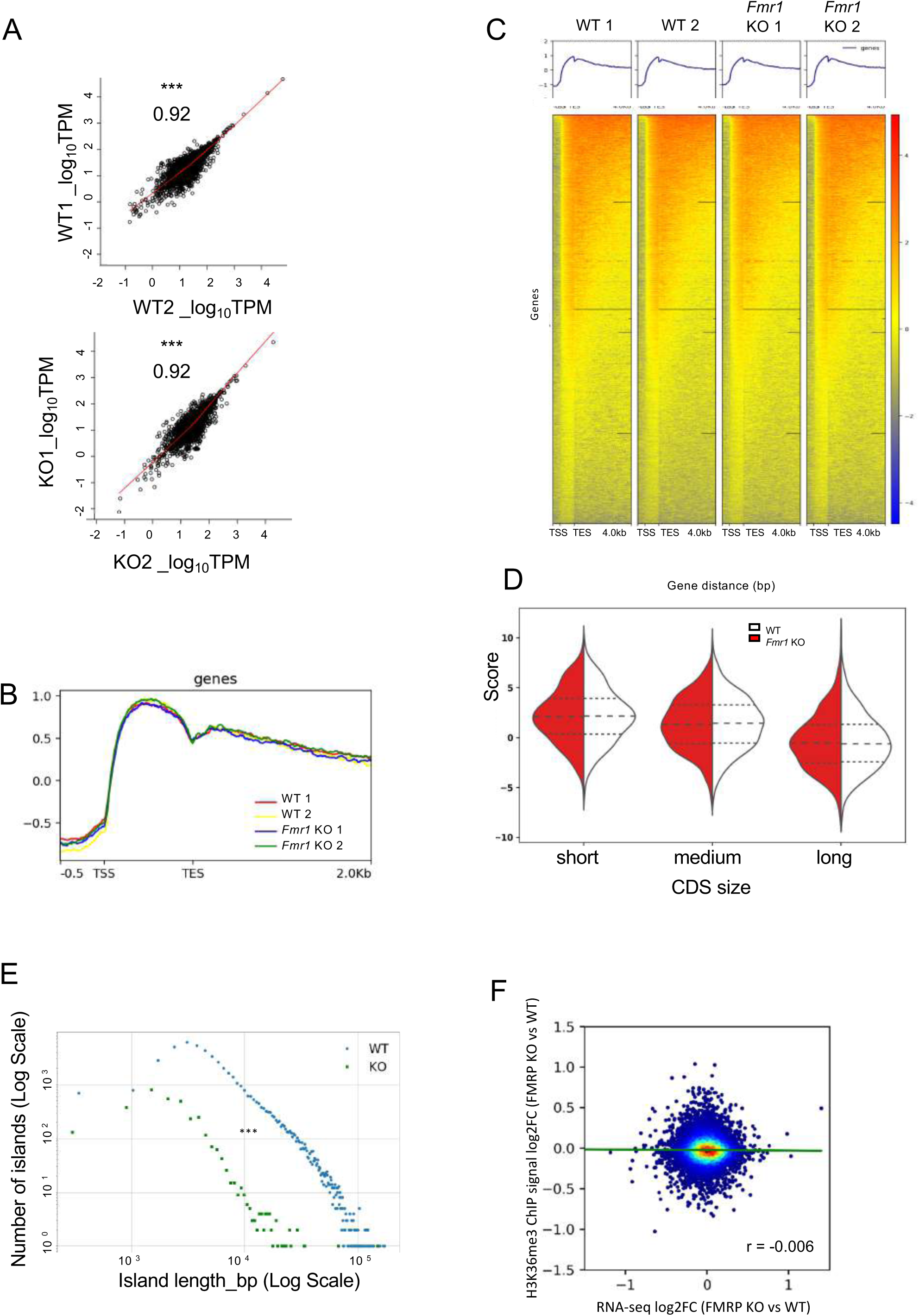
H3K36me3 ChIP metagene analysis (Related to Figure 6, also see Table S2) (A) Scatter plot for ChIP-seq normalized reads density (Transcripts per million mapped reads, TPM) mapped to the genic regions between biological replicate samples, WT1 and WT2 and *Fmr1* KO1 and *Fmr1* KO2. Pearson r values for correlation efficiency are shown. Asterisks indicate statistical significance. (B) Metagene plots using deepTools for distribution of H3K36me3 marks along the gene lengths. A similar increase in ChIP signal in all samples is seen in the gene body. The transcription start site (TSS), Transcription end site (TES) and 2.0kb downstream is shown. (C) Heatmap for distribution of H3K36me3 signal along the gene length in each sample, compiled plot in Figure S4B. The transcription start site (TSS), Transcription end site (TES) and 4.0kb downstream is shown. No overall differences in ChIP signal was observed between samples. (D) Violin plot to assess the effect of gene length on H3K36me3 ChIP signal between WT (white) and *Fmr1* KO (red) samples. (E) Histogram depicting number of islands (log scale) and their respective island length for all significant islands (bp in log scale) identified in the WT (blue) and *Fmr1* KO (green) ChIP-seq. The island lengths were parsed in bins of 100bp and the center of each bin is plotted in the histogram. Asterisks indicate statistical significance. (F) Scatter plot comparing the log2 fold changes in RNA-seq (x-axis) with the ChIP-seq data (y-axis) averaging H3K36me3 island differences within each gene is plotted. Pearson correlation coefficient value is stated.

**Figure S5.**
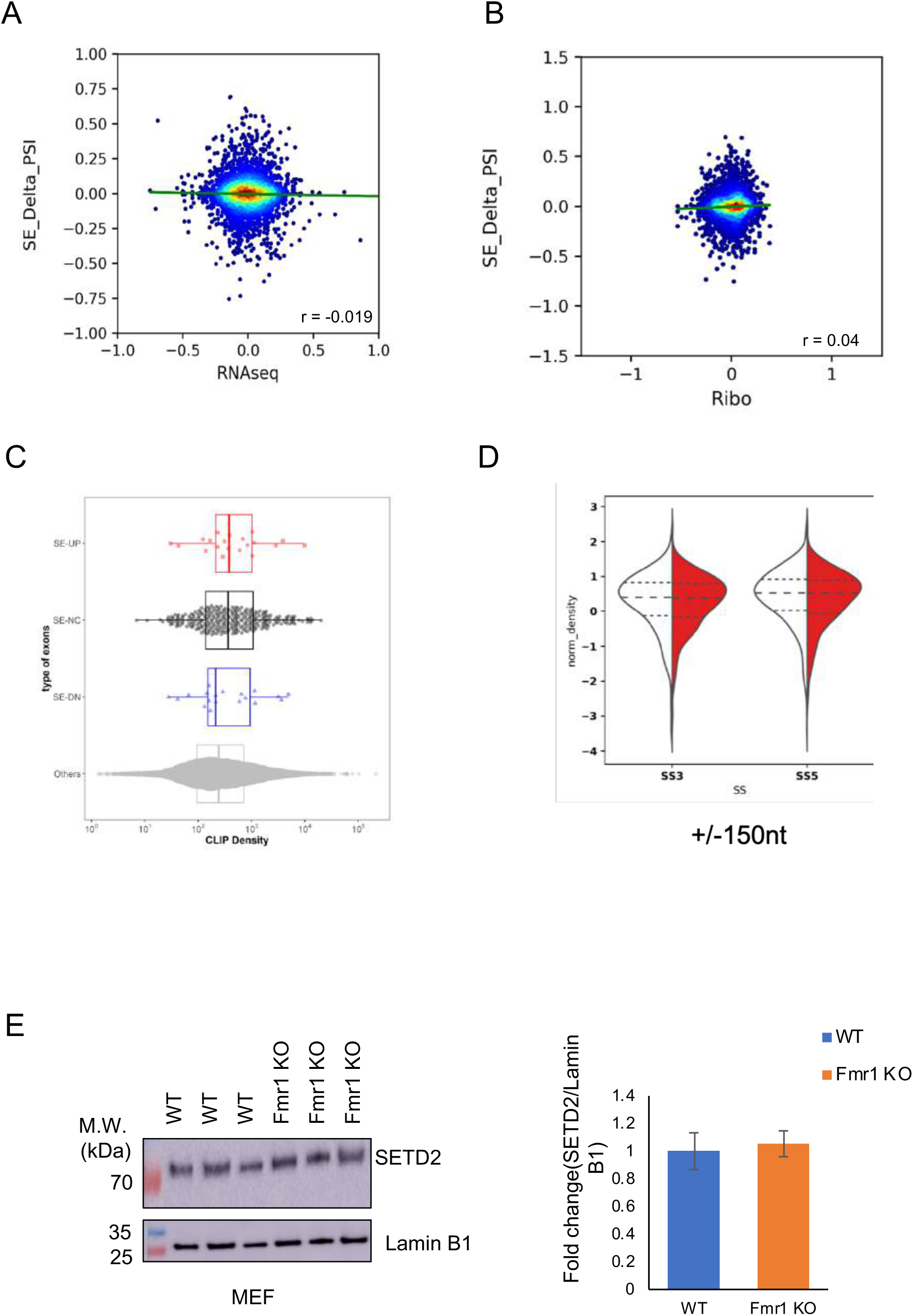
Comparison of skipped exon genes to RNA abundance and RPF levels (Related to Figure 1,6, and 7 also see Table S1 and S3) (A) Scatter plot for the delta percent spliced-in (Psi/Ψ) score of skipped exons (y-axis) (From Figure 7) versus their RNA fold changes (From Figure 1) on the x axis. Pearson correlation coefficient value is stated. (B) Scatter plot for the delta percent spliced-in (Psi/Ψ) score of skipped exons (y-axis) (From Figure 7) versus their RPF levels (From Figure 1) on the x axis. Pearson correlation coefficient value is stated. (C) Swarm plots are shown representing the FMRP CLIP density (CLIP level/length) using the FMRP CLIP-Seq data (GSE45148) at skipped exons and their flanking introns of genes with increased exon skipping (SE-UP in red), decrease skipped exons SE-DN in blue) and no significant change in skipped exons (SE-NC) are plotted. The other category represents exons with no alternative splicing detected. Wilcox test was used to determine significance of differences amongst the categories. (D) Violin plot for the H3K36me3 ChIP signal +/-150nt at the 5’ (SS5) and 3’ (SS3) splice sites of the alternatively skipped exons in WT (white) and *Fmr1* KO (red) hippocampus tissue (p-value<0.05, Wilcox test for significance). (E) Western blot analysis of SETD2 and lamin B1 in immortalized MEFs from WT and *Fmr1* KO mice (left), passage number 31. When quantified and made relative to Lamin B1 (right), there was no observable change in SETD2 levels in the KO (3 replicates, p=0.48, two-tailed t test). S.E.M is plotted (also see Figure 5)

